# Epigenetic modulation reveals differentiation state specificity of oncogene addiction

**DOI:** 10.1101/2020.05.04.077602

**Authors:** Mehwish Khaliq, Mohan Manikkam, Elisabeth D. Martinez, Mohammad Fallahi-Sichani

## Abstract

Hyperactivation of the MAPK signaling pathway motivates the clinical use of MAPK inhibitors for BRAF-mutant melanomas. Heterogeneity in differentiation state due to epigenetic plasticity, however, results in cell-to-cell variability in the state of MAPK dependency, diminishing the efficacy of MAPK inhibitors. To identify key regulators of such variability, we screened 276 epigenetic-modifying compounds, individually or combined with MAPK inhibitors, across genetically diverse and isogenic populations of melanoma cells. Following single-cell analysis and multivariate modeling, we identified three classes of epigenetic inhibitors that target distinct epigenetic states associated with either one of the lysine-specific histone demethylases KDM1A or KDM4B, or BET bromodomain proteins. While melanocytes remained insensitive, the anti-tumor efficacy of each inhibitor was predicted based on melanoma cells’ differentiation state and MAPK activity. Our systems pharmacology approach highlights a path toward identifying actionable epigenetic factors that extend the BRAF oncogene addiction paradigm on the basis of tumor cell differentiation state.

## Introduction

Therapeutic inhibition of oncogenic signaling often leads to variable responses due to cell-to-cell heterogeneity in the state of oncogene dependency ^1^. Heterogeneity may result from secondary genetic mutations. Alternatively, it may be caused by epigenetic differences associated with a cell’s developmental lineage or differentiation state ^2–10^. An example of such heterogeneity is observed among BRAF-mutated melanomas and causes fractional responses to BRAF/MEK-targeted therapies ^11–23^. Numerous studies have associated fluctuations in the state of MAPK dependency to melanoma differentiation state plasticity ^6,18,24–27^. Such plasticity spans a spectrum, ranging from a pigmented melanocytic phenotype associated with transcriptional regulators SOX10 and MITF ^28^, to a neural crest-like state that expresses NGFR ^12,29^, to an undifferentiated state characterized by high expression of receptor tyrosine kinases such as AXL ^19,30^. Melanoma tumors consist of a mixture of these phenotypes at variable single-cell frequencies ^25^. Although the consequences of such heterogeneities for drug resistance are widely recognized, there is still little known about actionable epigenetic factors that may link heterogeneity in the state of MAPK dependency to plasticity in differentiation state. A system-wide exploration of these factors may reveal novel opportunities for epigenetic treatments that will minimize the emergence of drug resistance. In principle, identifying epigenetic treatments that promote cellular requirement for MAPK signaling may enhance the efficacy of BRAF/MEK inhibitors when used in combination ^12,31–33^. Alternatively, treatments that target synthetic lethal partners of the BRAF oncogene may serve as a strategy to kill intrinsically drug-resistant cells ^34^.

In this paper, we took a systems pharmacology approach to test the hypothesis that heterogeneity in the state of MAPK dependency may result from a subset of key epigenetic variations across tumor cells of heterogeneous differentiation states. To identify key regulators of such variations, we used a library of 276 small molecule epigenetic modulators in BRAF-mutant melanoma cell lines that cover a wide spectrum of differentiation states. We evaluated the effect of each epigenetic modulator (individually and in combination with BRAF/MEK inhibitors) on cell survival, proliferation, differentiation state, and MAPK activity. Integrating multiplexed single-cell analysis with multivariate modeling and genetic experiments, we identified three classes of small molecules that target seemingly distinct epigenetic states in melanoma cells. These states included: (i) a lysine demethylase 1A (KDM1A)-dependent state, predominantly observed in undifferentiated cells, that is efficiently inhibitable by the reversible KDM1A inhibitor SP2509, (ii) a lysine demethylase 4B (KDM4B)- dependent state, associated with neural crest-like cells, that is sensitive to JIB-04 (a pan-inhibitor of Jumonji histone demethylases), and (iii) a state induced by BET bromodomain inhibitors such as OTX015 (Birabresib), which enhances cells’ requirement for MAPK signaling. Single-cell analysis shows that these states might co-exist in different combinations and frequencies, highlighting mutual epigenetic vulnerabilities among genetically diverse melanoma cell populations. Importantly, non-transformed primary melanocytes were not sensitive to these inhibitors. These results, therefore, provide a path for identifying actionable epigenetic factors that may extend the BRAF oncogene addiction paradigm on the basis of tumor cell differentiation state.

## Results

### Single-cell analysis uncovers heterogeneities in differentiation, proliferation and signaling states

To elucidate single-cell heterogeneities in differentiation state and their variations across melanoma cell populations, we utilized high-throughput, multiplex immunofluorescence microscopy. We exposed a panel of sixteen BRAF^V600E/D^ melanoma cell lines and a batch of non-transformed human primary epidermal melanocytes to the BRAF inhibitor vemurafenib (at 100 nM), alone or in combination with the MEK inhibitor trametinib (at 10 nM). For the purpose of comparison, we also included an NRAS^Q61K^-mutated variant of the A375 cell line, representing a common mechanism of acquired resistance to BRAF/MEK inhibitors ^35^. All cells were fixed following 3 to 5 days of treatment and protein levels of three validated differentiation state markers, MITF, NGFR and AXL (Supplementary Fig. 1), were quantified at a single-cell level (Extended Data Fig. 1, 2). To visualize baseline and treatment-induced variations in all three markers, we performed *t*-distributed stochastic neighbor embedding (t-SNE) analysis on a total population of 6069 randomly selected cells from all of the 102 tested conditions, covering the entire panel of cell lines, drugs and timepoints (Fig. 1a, Supplementary Fig. 2a, b). Single-cell analysis revealed a continuum of differentiation states ranging from MITF^High^ to NGFR^High^ to AXL^High^ cells. We then used the t-SNE map as a reference to visualize differentiation state variations in isogenic cell populations from each cell line (Fig. 1b, Supplementary Fig. 2c, d). We also quantified the extent of heterogeneity in each marker by computing the Fano factor, a standardized measure of dispersion of probability distribution (Supplementary Fig. 3). To assess the combined effect of heterogeneity in all three differentiation markers, we determined the average cell-to-cell distance in each cell population (Supplementary Fig. 4). All clonal cell lines exhibited a high degree of heterogeneity in at least one of the differentiation markers and most cell lines expressed substantial plasticity following exposure to BRAF/MEK inhibitors. Surprisingly, non-transformed melanocytes also exhibited heterogeneity at a level comparable to melanoma cell lines, suggesting that plasticity in differentiation state is not unique to cancer cells.

**Figure 1.**
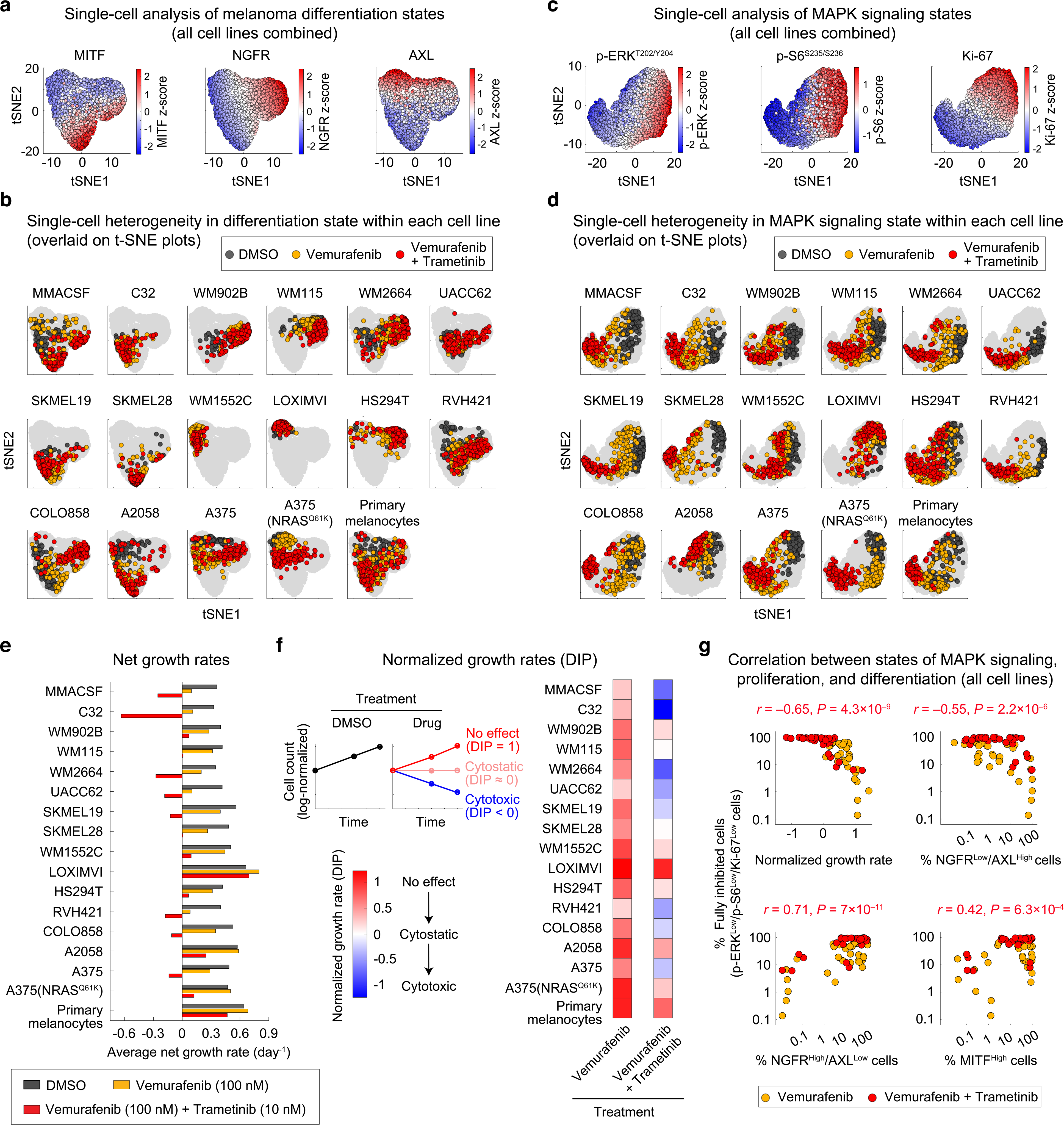
Single-cell analysis uncovers heterogeneities in melanoma differentiation, proliferation and MAPK signaling states across a wide range of BRAF/MEK inhibitor sensitivity. (a,b) Single-cell protein levels of three melanoma differentiation state markers, MITF, NGFR and AXL, measured by multiplexed immunofluorescence microscopy and overlaid on t-SNE plots. Cells were exposed to either vehicle (DMSO), BRAF inhibitor (vemurafenib at 100 nM), or the combination of BRAF and MEK inhibitors (vemurafenib at 100 nM and trametinib at 10 nM) for 3 and 5 days. Cells for each experimental condition were randomly selected from the pool of two replicates. Single-cell t-SNE maps for all cell lines combined (a) and projections of variation within each individual cell line (b) are shown. See also Supplementary Dataset 1. (c,d) Single-cell protein levels of p-ERK^T202/Y204^, p-S6^S235/S236^, and Ki67, measured by multiplexed immunofluorescence microscopy and overlaid on t-SNE plots. Treatment conditions are the same as in a, b. Single-cell t-SNE maps for all cell lines combined (c) and projections of variation within each individual cell line (d) are shown. See also Supplementary Dataset 2. (e) Average net growth rates calculated from measurements of live cell count (across at least two replicates) at three timepoints (including 0, 3, and 5 days) following exposure to DMSO, vemurafenib (at 100 nM) or vemurafenib (at 100 nM) plus trametinib (at 10 nM). See also Supplementary Dataset 3. (f) Drug-induced normalized growth rates (a.k.a. DIP rates) calculated by dividing the average net growth rate for drug-treated cells to that for DMSO-treated cells in each cell line. Normalized growth rates < 0 indicate a net cell loss (i.e. drug-induced cytotoxicity), a value of 0 represent no change in viable cell number (i.e. cytostasis), a value > 0 indicates a net cell gain, and a value of 1 represents no drug effect as cells grow at the same rate as in the DMSO condition. (g) Pairwise Pearson correlation analysis between the fraction of p-ERK^Low^/p-S6^Low^/Ki-67^Low^ cells (referred to as fully inhibited cells) and drug-induced normalized growth rate (top left), the fraction of undifferentiated (NGFR^Low^/AXL^High^) cells (top right), the fraction of neural crest-like (NGFR^High^/AXL^Low^) cells (bottom left), and the fraction of melanocytic (MITF^High^) cells (bottom right) across 16 melanoma cell lines treated with BRAF/MEK inhibitors for 3 and 5 days.

Because proliferation of BRAF-mutant cells is attributed to their aberrant MAPK signaling, we asked whether heterogeneity in differentiation state was associated with potentially distinct patterns in MAPK signaling. We thus multiplexed single-cell immunofluorescence measurements of p-ERK^T202/Y204^, Ki-67 (a proliferation marker), and p-S6^S235/S236^ (a marker of TORC1 activity that is upregulated in MAPK inhibitor-tolerant cells ^36^) in the same group of cell lines exposed to the same BRAF/MEK inhibitors for 3 to 5 days (Extended Data Fig. 3, 4). Fano factor, cell-to-cell distance and t-SNE analysis also revealed substantial variability in MAPK signaling (Fig. 1c, d, Supplementary Fig. 5, 6). While vemurafenib led to partial inhibition of MAPK signaling relative to drug-naïve cells, the combination of vemurafenib and trametinib suppressed the pathway more strongly and drove a larger proportion of cells toward a fully inhibited (p-ERK^Low^/p-S6^Low^/Ki-67^Low^) state (Fig. 1d). The fraction of fully inhibited cells, however, varied among cell lines. We hypothesized that such variability in MAPK signaling would explain differences in the overall MAPK inhibitor sensitivity and might be related to heterogeneity in melanoma differentiation state.

To compare the overall BRAF/MEK inhibitor sensitivity among cell lines, we computed drug-induced normalized growth rates (a.k.a DIP rates ^37^) by normalizing the average net growth rate of each cell line in the presence of each drug to that in DMSO-treated cells (Fig. 1e, f and Extended Data Fig. 5a). By correlating normalized growth rates to the state of MAPK signaling across 16 cell lines, we identified a strong correlation between the fraction of fully inhibited (p-ERK^Low^/p-S6^Low^/Ki-67^Low^) cells and the overall sensitivity to BRAF/MEK inhibitors (Pearson’s *r* = -0.65, *P* = 4.3×10^−9^) (Extended Data Fig. 5b and Fig. 1g; top left panel). We next extended the systematic correlation analysis to the diversity of differentiation states. We discovered that upon BRAF/MEK inhibition, predominantly undifferentiated (AXL^High^) cell lines generated the smallest populations of fully inhibited cells (Extended Data Fig. 5b and Fig. 1g; top right panel). BRAF/MEK inhibitor resistance in these cells was associated with a high frequency of proliferating (Ki-67^High^) cells exhibiting incomplete inhibition of the MAPK pathway. In contrast, populations of neural crest-like (NGFR^High^/AXL^Low^) cells or differentiated (MITF^High^) cells exhibited substantial co-inhibition of p-ERK, p-S6 and Ki-67 (Extended Data Fig. 5b and Fig. 1g; bottom panels). Drug adaptation in these populations was, therefore, associated with an overall reduced requirement for MAPK signaling.

Together, these analyses revealed a spectrum of heterogeneities in melanoma differentiation state that corresponded to two previously described but distinct ways through which cells may tolerate the effect of BRAF/MEK inhibition: (1) incomplete inhibition of the MAPK pathway (as seen in AXL^High^ cells), and (2) reduced requirement for MAPK signaling (as seen predominantly in NGFR^High^/AXL^Low^ cells).

### A chemical screen identifies epigenetic modulators of phenotypic heterogeneity

To systematically search for epigenetic factors that may link heterogeneity in melanoma differentiation state to MAPK dependency, we performed a multi-stage phenotypic screen using a library of 276 epigenetic-modifying compounds. Each compound was used to modulate either a key epigenetic writer, eraser or reader, or a related protein, after which phenotypic responses in the presence or absence of BRAF/MEK inhibitors were investigated. For the first stage of the screen, we selected COLO858 and MMACSF cell lines which show distinct patterns of differentiation states (Fig. 2a). Cells from both cell lines were initially exposed to two different doses (0.2 and 1 μM) of each of the 276 epigenetic compounds or vehicle (DMSO) for 24 h. Vemurafenib alone (at 100 nM), vemurafenib in combination with trametinib (at 10 nM), or vehicle (DMSO), was then added and cells were grown for a further 72 or 120 h prior to fixation (Fig. 2b and Supplementary Tables 1, 2). To differentiate the impact of epigenetic compounds, the growth rates for cells treated with each compound were compared to cells treated without any epigenetic treatment. Statistical analysis identified 58 compounds that led to a significant decrease in normalized growth rate in at least one of the tested conditions (Supplementary Fig. 7, 8).

**Figure 2.**
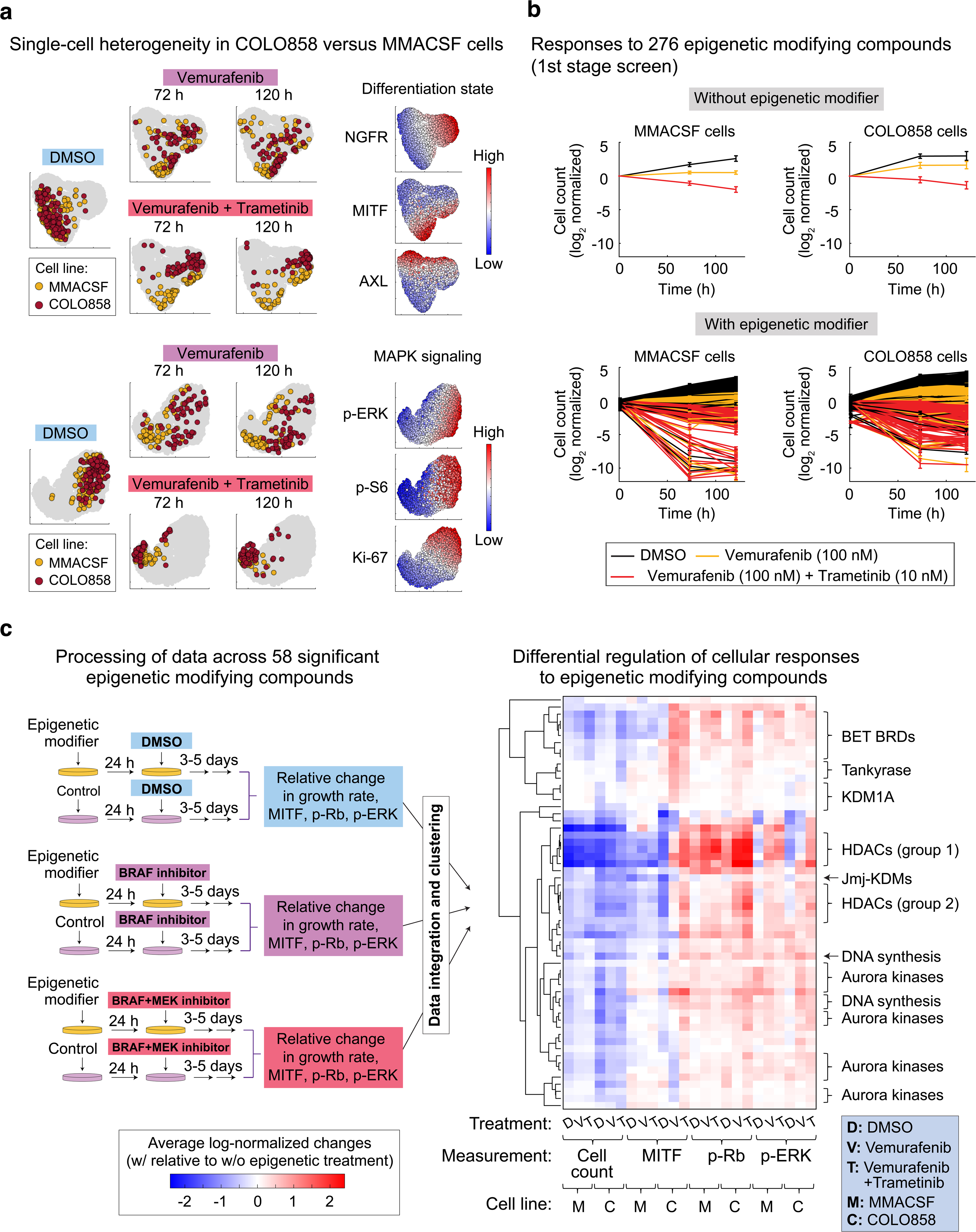
A chemical screen identifies epigenetic modulators of phenotypic heterogeneity. **(a)** t-SNE maps comparing single-cell heterogeneity in differentiation state (MITF, NGFR and AXL) and MAPK signaling (p-ERK^T202/Y204^, p-S6^S235/S236^ and Ki67) within two BRAF^V600E^ melanoma cell lines, MMACSF and COLO858, following exposure to vehicle (DMSO), vemurafenib (at 100 nM), alone or in combination with trametinib (at 10 nM), for 72 and 120 h. **(b)** Log_2_-normalized changes in live cell count following exposure of COLO858 and MMACSF cells to either DMSO, vemurafenib (at 100 nM), or vemurafenib (at 100 nM) plus trametinib (at 10 nM), for a period of 120 h. Cells were pretreated for 24 h with either DMSO (top panels) or two different doses (0.2 and 1 μM) of each of the 276 epigenetic-modifying compounds (bottom panels). Data represent mean values ± s.d. calculated across 2 replicates (in the presence of epigenetic treatments) and 276 replicates (in the absence of epigenetic of treatments). See also Supplementary Dataset 4. **(c)** A schematic representation of the overall procedure of data collection, processing (normalization), integration and hierarchical clustering using measurements of cellular grow rate, MITF, p-Rb^S807/811^, and p-ERK^T202/Y204^ at indicated treatment conditions and timepoints in MMACSF and COLO858 cells. Unsupervised clustering analysis was performed on data collected for 58 epigenetic compounds that led to a statistically significant decrease in normalized growth rate (when used either as a single agent, or in combination with BRAF/MEK inhibitors) in either or both of cell lines. Prior to clustering, data collected for cells treated with each epigenetic compound and MAPK inhibitor condition (i.e., DMSO, vemurafenib, or vemurafenib plus trametinib) were normalized to cells treated without any epigenetic compound and the same MAPK inhibitor condition. Groups of compounds with similar nominal epigenetic targets are listed on the right side. See also Supplementary Dataset 5.

To infer potential variations in the mechanisms of action of the epigenetic compounds, we co-stained cells for p-ERK^T202/Y204^, p-Rb^S807/811^, and MITF, quantifying changes induced by each of the 58 compounds in MAPK signaling, cell cycle progression, and differentiation state, respectively. Unsupervised clustering of the normalized responses revealed remarkable similarities among classes of compounds with common nominal epigenetic targets, suggesting that these responses are most likely the consequence of their on-target effects (Fig. 2c and Supplementary Fig. 9a). Each class of compounds induced responses that were either cell line-or MAPK inhibitor-specific, or common between both cell lines, or independent of MAPK inhibitor condition. For example, CUDC-907 (Fimepinostat), Quisinostat, Panobinostat, Dacinostat, and Trichostatin A were identified as a cluster of HDAC inhibitors (labeled as group 1 in Fig. 2c) that significantly reduced growth rate in both COLO858 and MMACSF cells to similar degrees and independent of the MAPK inhibitor conditions (Supplementary Fig. 9b). In contrast, KDM1A inhibitors inhibited net growth rate selectively in COLO858 cells, and they showed a higher efficacy in the absence of BRAF/MEK inhibitors.

### Correlated patterns of melanoma responses to mechanistically distinct epigenetic inhibitors

To identify potential relationships between the efficacy of each class of epigenetic compounds, we analyzed their effects across a more diverse group of cell lines. Thus, in the second stage of screen, we focused on seven epigenetic inhibitors representing the most effective classes (including HDAC inhibitors CUDC-907 and Givinostat, pan-Jmj-KDM inhibitor JIB-04, Tankyrase inhibitor AZ6102, BET inhibitors I-BET762 and OTX015, and KDM1A inhibitor SP2509) and a group of eight cell lines representing a wider spectrum of differentiation states (Fig. 3a, b). By testing each compound in non-transformed human primary melanocytes, we chose effective concentrations of each inhibitor that had little to no effect on healthy cells (Supplementary Fig. 10). To quantify the benefit resulting from combining each of the epigenetic inhibitors with vemurafenib and trametinib, we computed the deviation from Bliss independence (DBI), a metric that compares the observed cellular response to the combination treatment with that expected given independent action for the two individual treatments ^38^. We found that the HDAC inhibitor CUDC-907 induced substantial tumor cell killing in all of 8 melanoma cell lines and exhibited additive (independent) to synergistic responses when combined with vemurafenib and trametinib in 6 cell lines (Fig. 3b, c). Treatment with the other 6 epigenetic compounds uncovered more heterogeneous patterns of response across cell lines and MAPK inhibitor conditions. For example, BET inhibitors OTX015 and I-BET762 were not effective in any of the cell lines when used as a single agent, but they induced strong cytotoxic and synergistic responses in combination with vemurafenib and trametinib in the majority of cell lines. SP2509, on the other hand, was effective in 5 cell lines, while exhibiting antagonism in combination with vemurafenib and trametinib in all cell lines.

**Figure 3.**
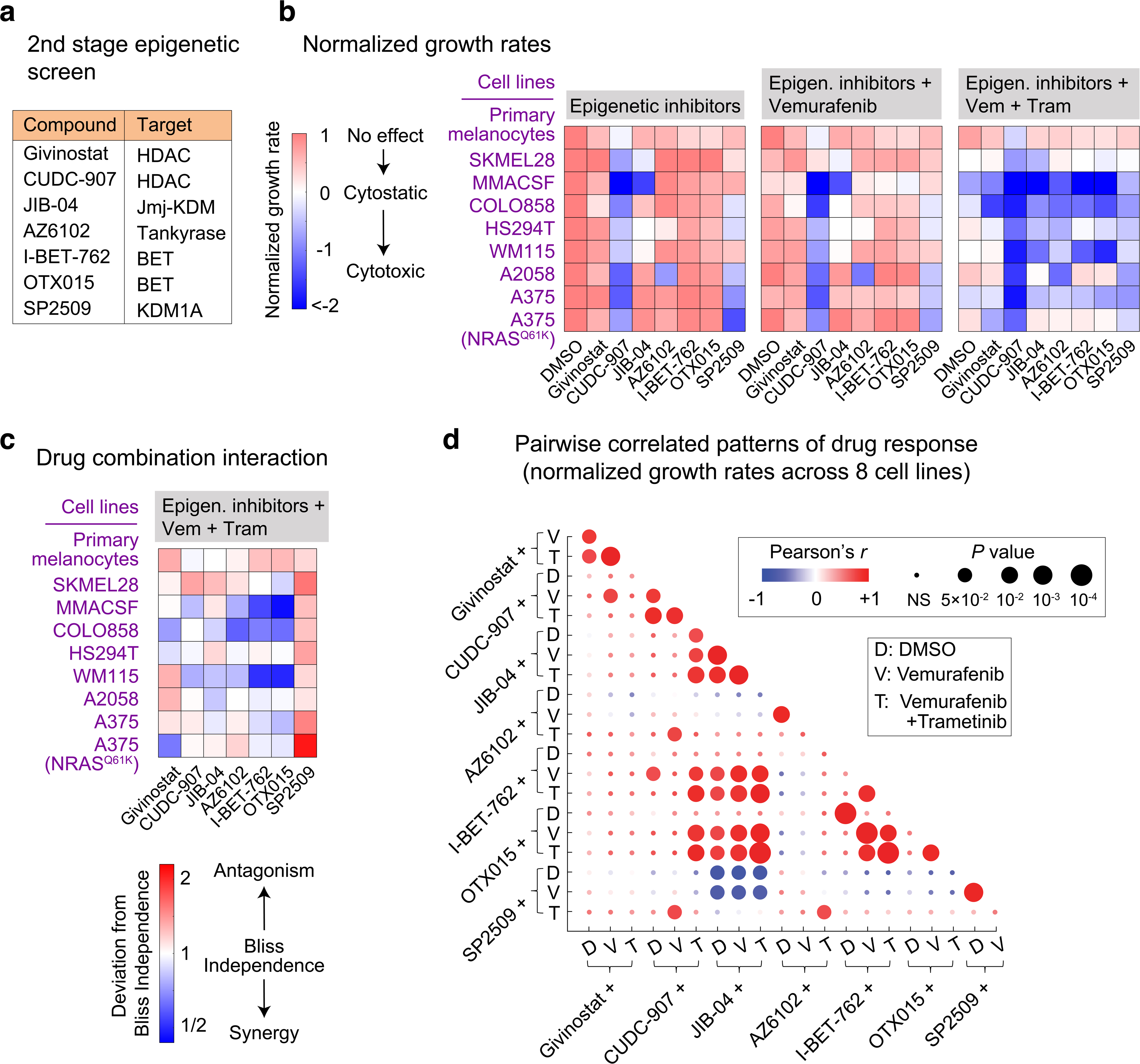
Patterns of melanoma responses to mechanistically distinct epigenetic inhibitors. **(a)** Epigenetic compounds (and their nominal protein targets) used in the second stage of epigenetic compound screen. **(b)** Measurements of normalized growth rate (averaged across at least 2 replicates) induced by each epigenetic inhibitor, when used either as a single agent (left), in combination with vemurafenib (middle), or in combination with vemurafenib plus trametinib (right), across eight different BRAF-mutant melanoma cell lines and non-transformed primary melanocytes. Treatment doses for different compounds are as follows: Givinostat (200 nM), CUDC-907 (20 nM), JIB-04 (200 nM), AZ6102 (1 μM), I-BET-762 (1 μM), OTX015 (0.2 μM), SP2509 (1 μM). Cells were pre-treated with the indicated epigenetic compounds for 24 h and then treated for a period of 3-5 days with either DMSO, vemurafenib (at 100 nM) or the combination of vemurafenib (at 100 nM) and trametinib (at 10 nM). See also Supplementary Dataset 6. **(c)** Deviation from Bliss Independence (DBI) values were computed across diverse epigenetic inhibitor treatments in combination with vemurafenib plus trametinib. DBI = 1 represents an independent (additive) effect equal to what is expected for the combination of drugs that act independently, DBI < 1 represents a combined effect stronger than expected for an independent combination (i.e. synergism), and DBI > 1 represents a combined effect weaker than expected for an independent combination (i.e. antagonism). **(d)** Pairwise Pearson correlations between the effects (i.e. normalized growth rates) of mechanistically distinct epigenetic inhibitors (used individually or in combination with vemurafenib or vemurafenib plus trametinib) evaluated across eight BRAF-mutant melanoma cell lines.

To systematically explore potential relationships between patterns of responses to epigenetic inhibitors, we computed all pairwise correlations between the efficacy of seven epigenetic compounds across the eight cell lines (Fig. 3d). As expected, the efficacy of two BET inhibitors, OTX015 and I-BET762, was strongly correlated (Pearson’s *r* = 0.99, *P* < 10^-4^). In addition, the efficacy of BET inhibitors and the pan-Jmj-KDM inhibitor JIB-04 was positively correlated when cells were co-treated with vemurafenib plus trametinib (*r* = 0.85, *P* < 10^-3^). In contrast, a negative correlation was observed between the efficacies of JIB-04 and SP2509. We thus asked whether such correlations would be maintained if these compounds were assayed across a larger panel of cell lines. By expanding the analysis to a total of sixteen cell lines, we confirmed the statistical significance of the negative correlation between responses to SP2509 and JIB-04 (Fig. 4a). In more than half of the cell lines tested, five days of treatment with SP2509 induced substantial cell killing (Fig. 4b and Extended Data Fig. 6a). SP2509 sensitivity correlated with sensitivity to SP2577 (seclidemstat; a clinical formulation of SP2509), which also suppressed melanoma cell growth (when used at a daily dose of 80 mg/kg) in corresponding melanoma xenografts (Extended Data Fig. 6b-d). Interestingly, however, cell lines with the highest level of resistance to SP2509 and SP2577 were sensitive to JIB-04, showing a range of responses from cytostatic to cytotoxic (Fig. 4a-c, Extended Data Fig. 6a). The triple combination of JIB-04, vemurafenib and trametinib led to additive (independent) to synergistic cell killing in most JIB-04-sensitive cell lines, whereas the combination of SP2509 with BRAF/MEK inhibitors was antagonistic in all SP2509-sensitive cell lines (Fig. 4c). When combined with vemurafenib and trametinib, both BET inhibitors OTX015 and I-BET762 also induced synergistic cell killing in most of the JIB-04-sensitive cell lines, while having minimal effects when used as a single agent. To determine if the epigenetic inhibitor effects persisted with long-term exposure, we extended the duration of growth assays. We found that the optimal epigenetic treatments identified for three of the MAPK inhibitor-resistant cell lines were highly efficacious, resulting in reduction of up to 1000-fold in live cell count over a period of 20 days (Fig. 4d).

**Figure 4.**
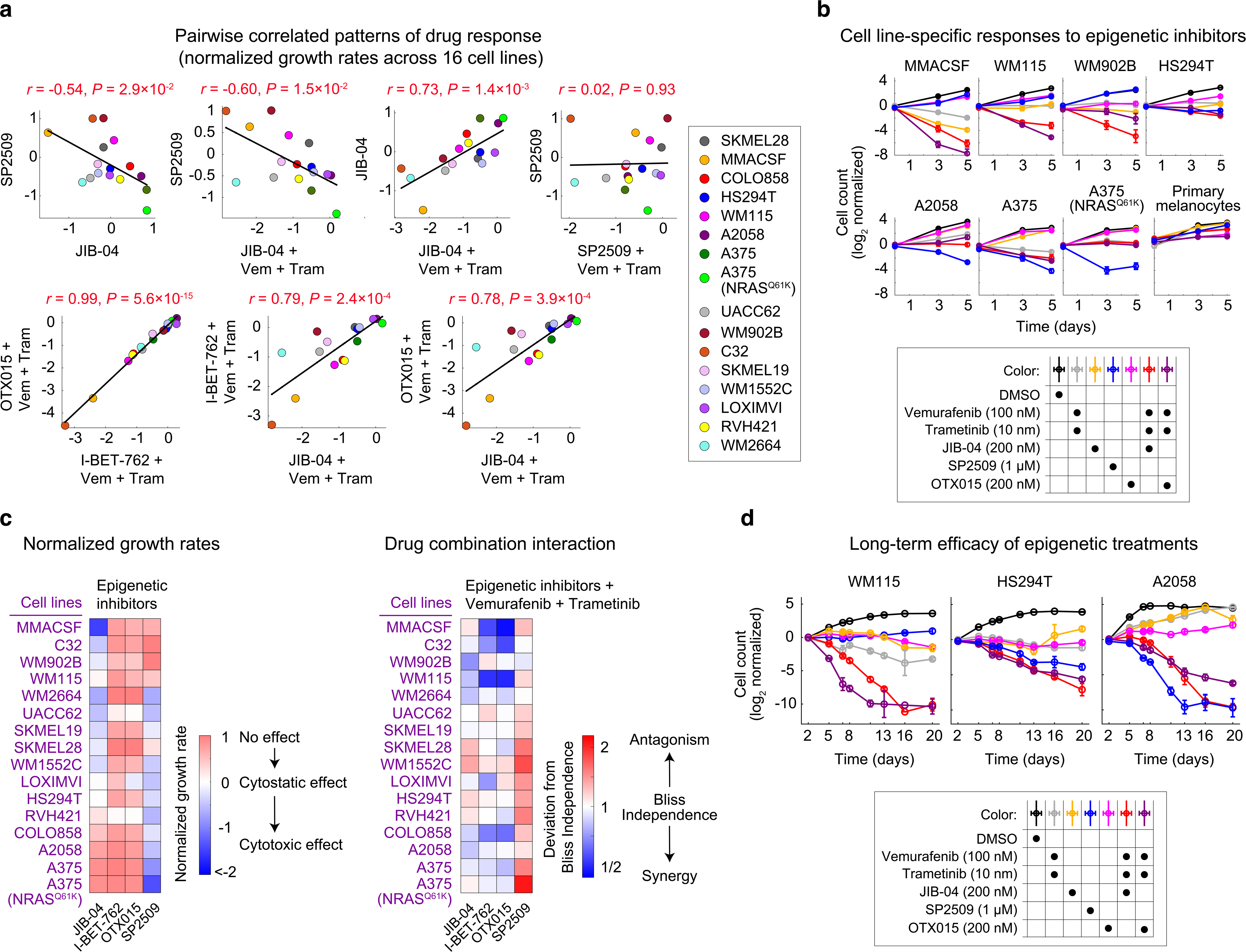
Correlated patterns of sensitivity to pharmacological inhibition of KDM1A, Jmj-KDMs and BET proteins across 16 BRAF-mutant melanoma cell lines. **(a)** Pairwise Pearson correlations between the effects (i.e. normalized growth rates) of KDM1A inhibitor SP2509 (at 1 μM), Jmj-KDM inhibitor JIB-04 (at 200 nM), and BET inhibitors OTX015 (at 200 nM) and I-BET762 (at 1 μM), used individually or in combination with vemurafenib (at 100 nM) plus trametinib (at 10 nM) and evaluated across sixteen BRAF-mutant melanoma cell lines. See also Supplementary Dataset 6. **(b)** Log_2_- normalized changes in live cell count following exposure of seven selected melanoma cell lines and non-transformed primary melanocytes to different drugs at indicated doses for a period of 5 days. MMACSF, WM115 and WM902B cell lines represent cell lines that exhibit high sensitivity to the combination of JIB-04 or BET inhibitors with BRAF/MEK inhibitors, while being resistant to SP2509. A2058, A375 and A375(NRAS^Q61K^) represent cell lines that are highly sensitive to SP2509. HS294T cells show partial sensitivity to either of the compounds. Data represent mean values ± s.d. calculated from two replicates per treatment condition. **(c)** Measurements of normalized growth rate (left) and deviation from Bliss independence (DBI) values computed across diverse epigenetic inhibitor treatments in combination with vemurafenib plus trametinib. Experimental conditions and the analysis approach are the same as in Fig. 3. **(d)** Log_2_-normalized changes in live cell count following exposure of three selected melanoma cell lines to different drugs at indicated doses for a period of 20 days. Data represent mean values ± s.d. across four replicates per treatment condition.

Together, our multi-stage epigenetic screen and systematic correlation analysis identified a set of seemingly distinct epigenetic states whose inhibition, either by the KDM1A inhibitor SP2509, or by the pan-Jmj-KDM inhibitor JIB-04 or BET inhibitors (when used in combination with BRAF/MEK inhibitors), led to tumor cell killing in a melanoma cell line-specific manner. Next, we asked whether such cell line-specific patterns of response may be linked to heterogeneity in MAPK signaling, proliferation and differentiation states.

### Multivariate modeling identifies predictors of epigenetic inhibitor efficacy

To determine whether molecular markers of differentiation state, MAPK activity or other phenotypic markers could predict the differential efficacy of epigenetic treatments, we measured both baseline levels and treatment-induced variations in MITF, NGFR, AXL, SOX10, p-ERK^T202/Y204^, p-S6^S235/S236^ and Ki-67 across eight melanoma cell lines. Given the possible contribution of epigenetic histone modifications in DNA damage repair ^39^, we also included phosphorylated Histone p-H2A.X^S139^ (a marker of DNA damage response) in our analysis. Multiplexed immunofluorescence measurements revealed how multiple protein markers were up- or downregulated depending on the cell line, MAPK inhibitor condition and epigenetic treatment (Fig. 5a). We then used partial least square regression (PLSR) analysis ^40^ to generate models that linked epigenetic treatment-induced changes in growth rates (“response” variables) to “input” vectors that combined baseline and treatment-induced changes in signaling and phenotypic data. Models were evaluated using leave-one-out cross-validation (Extended Data Fig. 7a). Overall, PLSR models built for JIB-04, SP2509, I-BET762 and OTX015 proved remarkably accurate with an average Pearson’s correlation coefficient of 0.85 ± 0.06 between the measured and predicted responses (Fig. 5b and Extended Data Fig. 7b).

**Figure 5.**
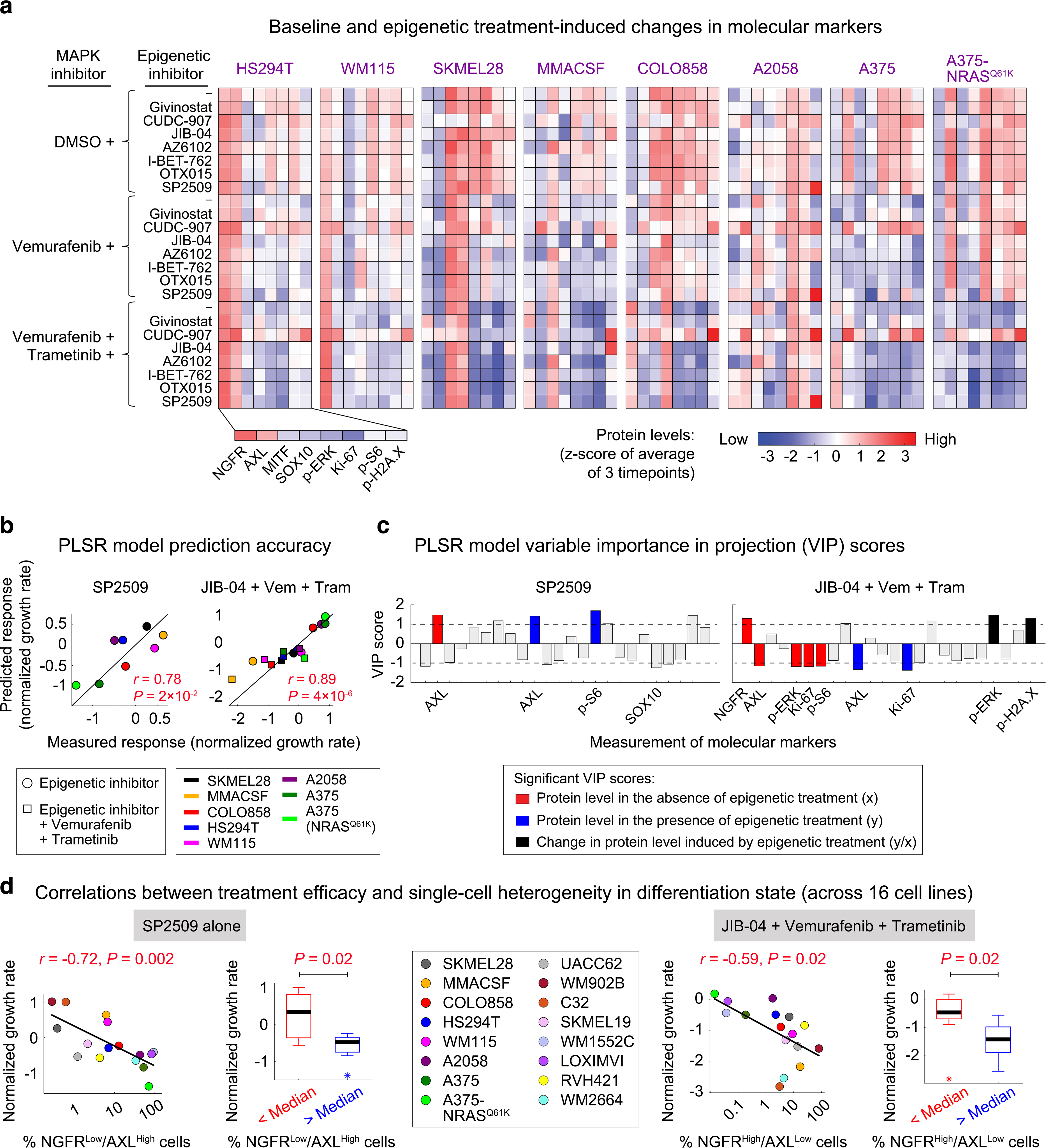
Multivariate modeling identifies differentiation state-specific predictors of epigenetic inhibitor efficacy. **(a)** Baseline and treatment-induced changes in protein measurements of melanoma differentiation state markers (NGFR, AXL, MITF, SOX10), MAPK signaling protein modifications (p-ERK^T202/Y204^, p-S6^S235/S236^), a proliferation marker (Ki-67), and a DNA damage response marker (p-H2A.X^S139^), across eight melanoma cell lines following treatment with the indicated epigenetic compounds, either individually or in combination with vemurafenib, or vemurafenib plus trametinib. Treatment doses for different compounds are as follows: Givinostat (200 nM), CUDC-907 (20 nM), JIB-04 (200 nM), AZ6102 (1 μM), I-BET-762 (1 μM), OTX015 (0.2 μM), SP2509 (1 μM), vemurafenib (100 nM), trametinib (10 nM). Protein data shown for each condition represent mean values across three timepoints and two replicates and are then z-scored across all cell lines and treatment conditions. See also Supplementary Datasets 7-10. **(b)** Pairwise Pearson correlation between responses (normalized growth rates) to SP2509 (left) or JIB-04 in combination with vemurafenib and trametinib (right), measured for each of the eight melanoma cell lines (x-axis) and corresponding responses predicted by partial least square regression (PLSR) modeling following leave-one-out cross validation (y-axis). **(c)** PLSR-derived variable importance in the projection (VIP) scores, highlighting combinations of protein measurements at the baseline (shown in red), following epigenetic inhibitor treatment (shown in blue), or the ratio of change induced by each epigenetic compound (shown in black), that are predictive of efficacy for SP2509 (left) and JIB-04 in combination with vemurafenib and trametinib (right). The sign of VIP score shows whether the change in variable correlated negatively or positively with the treatment-induced response. Only VIP scores of greater than 1 or smaller than -1 with a statistically significant Pearson correlation (*P* < 0.05) are highlighted. **(d)** Pairwise Pearson correlation analysis between the baseline fractions of NGFR^Low^/AXL^High^ cells and measurements of normalized growth rate in response to SP2509 when used as a single agent (left), and between the baseline fractions of NGFR^High^/AXL^Low^ cells and response to JIB-04 in combination with vemurafenib and trametinib (right), across sixteen BRAF-mutant melanoma cell lines. The significance of differences between normalized growth rates was also evaluated based on nonparametric Mann-Whitney U-test. For this test, sixteen cell lines were divided into two groups of eight based on whether the measured variable had a value above or below the median. Data for each group were then presented using box-and-whisker plots and median parameter values and interquartile ranges; bars extending to 1.5× the interquartile range are shown for each condition as a measure of variance. Parameter values for outlier cell lines are marked with asterisks.

To identify those measurements that are predictive of treatment efficacy, we computed the Variable Importance in the Projection (VIP) scores for each PLSR model ^41^ (Fig. 5c and Extended Data Fig. 7c). Among the most important determinants of treatment efficacy were differentiation markers AXL and NGFR as well as measurements of MAPK activity, including p-ERK, p-S6 and Ki-67. Informed by the PLSR results, single-cell analysis across the entire panel of cell lines showed that SP2509 was most effective in inhibiting NGFR^Low^/AXL^High^ populations of cells (Fig. 5d; left panels). In contrast, NGFR^High^/AXL^Low^ populations were sensitive to the triple combination of JIB-04, vemurafenib and trametinib (Fig. 5d; right panels). Additionally, BET inhibitors I-BET762 and OTX015 (when combined with BRAF/MEK inhibitors) enhanced tumor cell killing most significantly in populations of MITF^Low^/AXL^Low^ cells (Extended Data Fig. 7d). Surprisingly, global changes in differentiation state markers induced by 5 days of treatment with neither of the epigenetic compounds were identified as statistically significant by PLSR models. Only for SP2509-treated cells, single-cell analysis showed partial down-regulation of AXL and up-regulation of MITF in SP2509-sensitive cell lines (Supplementary Fig. 11).

Together, these data reveal how correlated patterns of responses to SP2509, JIB-04 and BET inhibitors are linked to the state of MAPK signaling and differentiation state. Cells in undifferentiated and neural crest-like states represent two different forms of MAPK inhibitor tolerance observed at variable frequencies across most melanoma tumors. Their selective sensitivity to the identified epigenetic inhibitors, therefore, supports the hypothesis that there are epigenetic features associated with melanoma differentiation state that may be linked to their state of BRAF/MAPK dependency.

### KDM4B and ZNF217 protein levels predict differentiation state-specific sensitivity to JIB-04 and SP2509

We sought to utilize our knowledge of interactions between JIB-04 and SP2509 and their protein targets to better understand the origins of their selective efficacy in melanoma cells. JIB-04 is known as a pan-inhibitor of Jmj-KDMs, suppressing the activities of KDM4A, KDM4B, KDM5A and KDM5B, all with IC_50_’s < 0.5 μM ^42^. We thus asked which of the Jmj-KDM proteins targeted by JIB-04 might explain its inhibitory effect on melanoma cells. We first depleted each of KDM4A, KDM4B, KDM5A and KDM5B proteins in two JIB-04-sensitive cell lines (WM115 and WM902B) and one JIB-04-resistant cell line (A2058) using four target-specific siRNAs combined into a single pool (Extended Data Fig. 8). Only depletion of KDM4B in WM115 and WM902B cells led to a statistically significant decrease in live cell count and enhanced cells’ sensitivity to the combination of vemurafenib and trametinib (Fig. 6a). To rule out the possibility of off-targeting by KDM4B siRNAs, we then examined the effects of three constituent KDM4B siRNAs independently. All individual siRNAs against KDM4B reduced KDM4B expression and cell viability in both WM115 and WM902B, while having no impact on KDM4A expression in these cell lines (Extended Data Fig. 9a, b). A2508 cells, on the other hand, were sensitive to KDM1A knockout by CRISPR using three independent sgRNAs (Fig. 6b and Extended Data Fig. 9c), which correlated with their sensitivity to SP2509.

**Figure 6.**
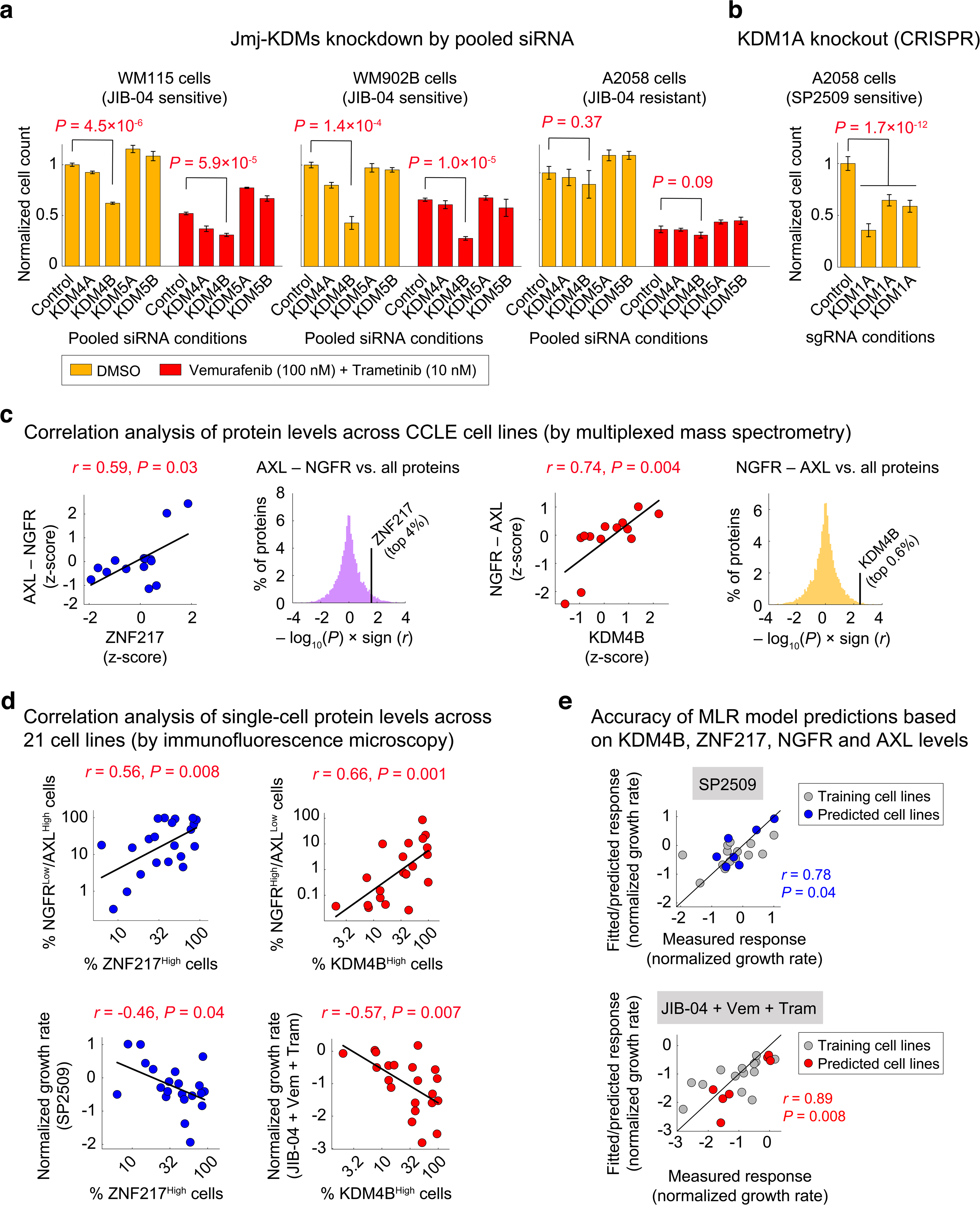
KDM4B and ZNF217 protein levels predict differentiation state-specific sensitivity to JIB-04 and SP2509. **(a)** Relative cell viability in WM115 cells (left), WM902B cells (middle) and A2058 cells (right) following treatment with indicated doses of the combination of vemurafenib and trametinib or vehicle (DMSO) in the presence of pools of four siRNAs targeting either KDM4A, KDM4B, KDM5A, and KDM5B for 96 h. Viability data for each treatment condition were normalized to cells treated with DMSO and the non-targeting (control) siRNA. Data represent mean values ± s.d. calculated across 3 replicates. Statistical significance was determined by two-sided *t* test. **(b)** Relative cell viability in Cas9-positive A2058 cells following treatment with three different types of KDM1A lentiviral single guide RNA (sgRNA) for 96 h. Viability data were normalized to cells treated with non-targeting (control) sgRNA. Data represent mean values ± s.d. calculated across 4 replicates. Statistical significance was determined by two-sided *t* test. **(c)** Pairwise Pearson correlation between variations in the difference between NGFR and AXL protein levels and ZNF217 (left panels) or KDM4B (right panels) in comparison with the rest of the measured proteome (shown as histograms) across BRAF-mutant melanoma cell lines. Protein data are extracted from the Cancer Cell Line Encyclopedia (CCLE) proteomics database (measured by multiplexed mass spectrometry) and z-scored across all of BRAF-mutant melanoma cell lines present in the database. **(d)** Pairwise Pearson correlation between the baseline fractions of ZNF217^High^ cells and KDM4B^High^ cells and the fractions of NGFR^Low^/AXL^High^ cells (top left), the fractions of NGFR^High^/AXL^Low^ cells (top right), sensitivity to SP2509 at 1 μM (bottom left), and sensitivity to the combination of JIB-04 (at 0.2 μM), vemurafenib (at 100 nM) and trametinib (at 10 nM) (bottom right), as evaluated by normalized growth rates following 5 days of treatment. Protein data were measured by immunofluorescence microscopy, quantified following single-cell analysis, and averaged across 3 replicates. See also Supplementary Datasets 11-14. **(e)** Pairwise Pearson correlation between responses (normalized growth rates) to SP2509 (top) or JIB-04 in combination with vemurafenib and trametinib (bottom), measured for each of the 21 melanoma cell lines (x-axis) and corresponding responses fitted by multi-linear regression (MLR) analysis of 14 cell lines (training set; shown in gray) or responses predicted by the trained models for the remaining 7 cell lines (validation set; shown in blue or red).

We then asked whether selective efficacy of SP2509 and JIB-04 might be explained by the differentiation state-specific expression of their epigenetic targets. We used mass spectrometry-based proteomics data for BRAF-mutant melanoma lines in the Cancer Cell Line Encyclopedia (CCLE) ^43^ to evaluate possible correlations between the expression of each epigenetic target and the relative expression of NGFR and AXL. Since SP2509 acts by blocking KDM1A interaction with its coactivator ZNF217 ^44^, we also included this protein in our analysis. While KDM1A protein levels did not significantly correlate with melanoma differentiation state (Supplementary Fig. 12a), ZNF217 was among the top 4% proteins that correlated with the expression of AXL relative to NGFR (Fig. 6c; left panels). In addition, levels of KDM4B (but not other JIB-04 targets) were among the top 0.6% proteins whose expression was significantly greater in melanoma cell lines that expressed higher levels of NGFR relative to AXL (Fig. 6c; right panels, and Supplementary Fig. 12b). These data suggest that the relative levels of KDM4B and ZNF217 proteins and differentiation markers NGFR and AXL may predict the selective sensitivity of melanoma cells to JIB-04 and SP2509.

To independently test this hypothesis, we first profiled the levels of all four proteins across our original panel of 16 melanoma cell lines plus an additional group of 5 cell lines by immunofluorescence microscopy (Extended Data Fig. 10a-d). Following single-cell analysis, we performed correlation analyses similar to those performed using the mass spectrometry data from CCLE. Our analysis confirmed significant pairwise correlations between the fractions of NGFR^Low^/AXL^High^ cells and ZNF217^High^ cells on one hand and the fractions of NGFR^High^/AXL^Low^ cells and KDM4B^High^ cells on the other hand (Fig. 6d; top panels). As expected, the sensitivity of melanoma cell lines to SP2509 and the combination of JIB-04, vemurafenib and trametinib were also correlated with the fractions of ZNF217^High^ and KDM4B^High^ cells, respectively (Fig. 6d; bottom panels). To further validate the predictivity of these protein markers, we divided the panel of 21 cell lines into a test set of 14 cell lines and a validation set of 7 cell lines. We then used the training set to develop multi-linear regression (MLR) models of treatment efficacy for SP2509 and JIB-04 using protein levels of NGFR, AXL, KDM4B and ZNF217. The trained models predicted SP2509 and JIB-04 efficacy in the validation set with high accuracy (Fig. 6e, Extended Data Fig. 10e). Together, these data demonstrate that the sensitivity of BRAF-mutant melanoma cells to SP2509 and JIB-04 can be predicted based on their differentiation state (NGFR versus AXL) and the relative levels of ZNF217 and KDM4B proteins.

## Discussion

Lineage-specific epigenetic mechanisms and their reprograming following oncogene inhibition can generate drug-tolerant states that diminish the efficacy of targeted therapies ^45^. In BRAF-mutant melanomas, comparable states tolerant of MAPK inhibition have been associated with variations in differentiation state. Through high-throughput profiling of human melanoma cell lines, we uncovered recurrent patterns of differentiation state heterogeneity that are comparable to those previously seen in patient tumors ^25^. We found that a melanoma cell’s ability to tolerate BRAF/MEK inhibitors was associated with its differentiation state. Responses of undifferentiated cells were associated with incomplete inhibition of the MAPK pathway, whereas MAPK-inhibited neural crest-like cells adapted to treatment by reducing their requirement for MAPK signaling.

To identify epigenetic features linked to melanoma differentiation states and MAPK dependency, we performed an epigenetic compound screen and identified three classes of compounds that target distinct melanoma cell states associated with either one of the lysine-specific histone demethylases KDM1A or KDM4B, or BET proteins. While non-transformed melanocytes remained unaffected by these compounds, their efficacy in melanoma cells depended on a cell’s state of MAPK activity and differentiation state. The KDM1A inhibitor SP2509 was most effective in inhibiting undifferentiated populations of cells that were intrinsically insensitive to BRAF/MEK inhibitors, whereas the combination of SP2509 with BRAF/MEK inhibitors led to antagonistic interactions. This is consistent with a recent finding regarding the role that KDM1A plays in disabling BRAF^V600E^ oncogene-induced senescence ^34^. KDM1A inhibition may, therefore, require MAPK signaling to restore senescence in NGFR^Low^/AXL^High^ cells. In contrast, NGFR^High^/AXL^Low^ populations of cells exhibited additive to synergistic responses to the combination of KDM4B and BRAF/MEK inhibition. BET inhibitors had minimal effect on melanoma cells when used as a single agent but led to tumor cell killing when combined with BRAF/MEK inhibitors. Our systematic studies, therefore, extend previous findings about the potential of each of the epigenetic regulators, KDM1A, Jmj-KDMs and BET proteins, as therapeutic vulnerabilities in melanomas ^20,34,46^ in two complementary ways. First, by associating these vulnerabilities to the patterns of single-cell heterogeneity in differentiation state, we explain how they may vary from one tumor to another. Second, by linking epigenetic vulnerabilities to the state of BRAF/MAPK dependency, we determine whether each epigenetic inhibitor reaches its maximal efficacy when used as a single agent or when combined with BRAF/MEK inhibitors. Future work should determine the specific mechanisms through which inhibition of each regulator might induce melanoma cell killing.

Cell-to-cell heterogeneity in the state of oncogene dependency poses a general challenge to the use of cancer targeted therapies. Our systems pharmacology approach provides a promising avenue toward the identification of actionable epigenetic factors that may extend the oncogene addiction paradigm on the basis of tumor cell differentiation state. We demonstrate the utility of this approach in BRAF-mutant melanomas by using JIB-04 and SP2509, as investigational tools, to block epigenetically diverse populations of MAPK inhibitor-tolerant cells. We show that the relative protein levels of KDM4B and ZNF217 (a KDM1A coactivator), determine differentiation state-specific sensitivity of melanoma cells to their corresponding inhibitors. Future work may gauge the mechanistic relationship between the activity of these proteins and melanoma differentiation state at a single-cell level.

## Methods

### Cell lines and reagents

BRAF-mutant melanoma cell lines used in this study were obtained from the following sources: COLO858 (from ECACC), RVH421 (from DSMZ), A375, A375(NRAS^Q61K^), C32, A2058, WM115, SKMEL28, HS294T, WM1552C, HS695T, RPMI7951, SKMEL5, A101D, IGR39, and human adult primary epidermal melanocytes (all from ATCC), LOXIMV1 (from DCTD Tumor Repository, National Cancer Institute), MMACSF (RIKEN BioResource Center), WM902B and WM2664 (from Wistar Institute), UACC62 and SKMEL19 (from the Cancer Cell Line Encyclopedia). All of the cell lines have been periodically subjected to re-confirmation by Short Tandem Repeat (STR) profiling by ATCC and mycoplasma testing by MycoAlert^TM^ PLUS mycoplasma detection Kit (Lonza). A375, A375(NRAS^Q61K^), A2058, HS294T, A101D and IGR39 cells were grown in DMEM with 4.5 g/l glucose (Corning, Cat# 10-013-CV) supplemented with 5% fetal bovine serum (FBS). RPMI7951, SKMEL5 and HS695T cells were grown in EMEM (Corning, Cat# 10-009-CV) supplemented with 5% FBS. C32, MMACSF, SKMEL28 and WM115 cells were grown in DMEM/F12 (Gibco, Cat# 11330-032) supplemented with 1% Sodium Pyruvate (Invitrogen) and 5% FBS. COLO858, LOXIMVI, RVH421, SKMEL19, UACC62, WM1552C and WM902B cells were grown in RPMI 1640 (Corning, Cat# 10-040-CV) supplemented with 1% Sodium Pyruvate and 5% FBS. Primary epidermal melanocytes were grown in Dermal Cell Basal Medium (ATCC, Cat# PCS-200-030) supplemented with Adult Melanocyte Growth Kit (ATCC, Cat# PCS-200-042). We added penicillin (50 U/ml) and streptomycin (50 lg/ml) to all growth media.

Small molecule inhibitors, including a library of 276 epigenetic-modifying compounds, chemicals used in the follow-up cell-based assays (Givinostat, CUDC-907, JIB-04, AZ6102, I-BET-762, OTX015 and SP2509), as well as vemurafenib and trametinib were all purchased from Selleck Chemicals. SP2577 was purchased from MedChem Express. The complete list of compounds used in this study, their catalog numbers and purity, as evaluated by HPLC and MS analysis, are presented in Supplementary Table 1. Compounds used for cell-based studies were dissolved in the appropriate vehicle (either DMSO or water) at a stock concentration of 10 mM. The vehicle for SP2577 used for *in vivo* studies is described below.

The following primary monoclonal antibodies (mAb, clone name) and polyclonal antibodies (pAb) with specified animal sources, catalog numbers and dilution ratios, were used in immunofluorescence staining assays: MITF (mouse mAb, clone D5, Abcam, Cat# ab3201, 1:800), p-ERK^T202/Y204^ (rabbit mAb, clone D13.14.4E, Cell Signaling Technology, Cat# 4370, 1:800), Ki-67 (mouse mAb, clone 8D5, Cell Signaling Technology, Cat#9449, 1:1200), AXL (goat pAb, R&D Systems, Cat# AF154, 1:400), p-Rb^S807/811^ (goat pAb, Santa Cruz Biotechnology, Cat# sc-16670, 1:400), NGFR (rabbit mAb, clone D4B3, Cell Signaling Technology, Cat# 8238, 1:1600), p-S6^S235/S236^ (rabbit mAb, clone D57.2.2E, Cell Signaling Technology, Cat# 4851, 1:400), SOX10 (mouse mAb, clone SOX10/991, Abcam, Cat# ab212843, 1:1200), p-H2A.X^S139^ (rabbit mAb, clone EP854(2)Y, Abcam, Cat# ab195188, 1:700), KDM1A (rabbit mAb, clone C69G12, Cell Signaling Technology, Cat# 2184, 1:1600), KDM4A (rabbit mAb, clone C37E5, Cell Signaling Technology, Cat# 5328, 1:100), KDM4B (rabbit mAb, clone D7E6, Cell Signaling Technology, Cat# 8639, 1:100), KDM5A (rabbit mAb, clone EPR18651, Abcam, Cat# ab194286, 1:1600), KDM5B (rabbit mAb, clone EPR12704, Abcam, Cat# ab181089, 1:100), and ZNF217 (rabbit pAb, Thermo Fisher Scientific, Cat# 720352, 1:200). The following secondary antibodies with specified sources and catalog numbers were used at a 1:2000 dilution: anti-rabbit Alexa Fluor 488 (Thermo Fisher, Cat# A21206), anti-mouse Alexa Fluor 647 (Thermo Fisher, Cat# A31571), anti-goat Alexa Fluor 568 (Thermo Fisher, Cat# A11057), anti-mouse Alexa Fluor 568 (Thermo Fisher, Cat# A10037), and anti-rabbit Alexa Fluor 647 (Thermo Fisher, Cat# A31573).

### Immunofluorescence staining, quantitation and analysis

Cells in 96-well plates were fixed in either 4% paraformaldehyde (PFA) for 20 min at room temperature or 100% ice-cold methanol for 15 min at -20°C. Cells were then washed with PBS, permeabilized in methanol (in cases where they were fixed with 4% PFA) for 10 min at -20°C, rewashed with PBS, and blocked using Odyssey blocking buffer (LI-COR Biosciences) for 1 h at room temperature. Cells were incubated overnight (∼16 h) at 4°C with primary antibodies in Odyssey blocking buffer. The following day, cells were washed three times with PBS supplemented with 0.1% Tween-20 (Sigma-Aldrich) (PBS-T) and incubated for 1 h at room temperature with the secondary rabbit, goat, or mouse antibodies. Cells were then washed twice with PBS-T, once in PBS, and then incubated with Hoechst 33342 (Thermo Fisher, Cat# H3570, 1:20000) for 20 min at room temperature. Cells were washed twice with PBS and imaged with a 10x objective using the ImageXpress Micro Confocal High-Content Imaging System (Molecular Devices) or the Operetta CLS High-Content Imaging System (Perkin Elmer). A total of 9 sites were imaged per well. Background subtraction was performed with ImageJ. Image segmentation and quantification of signal intensities in the whole cell, nucleus, cytoplasm, or the nucleus/cytoplasm (N/C) ratio were performed with CellProfiler ^47^. Population-average and single-cell data were analyzed using MATLAB 2019b. By generating histograms of single-cell data across a variety of conditions for each protein (X), including positive and negative controls, we identified an appropriate binary gate, based on which the percentage of X^High^ versus X^Low^ cells in each condition was quantified.

### Single-cell analysis and dimensionality reduction

To visualize single-cell heterogeneities in a two-dimensional space, we used a population of 6069 cells assembled following random selection of 60 individual cells (or fewer in the case of the highly drug-sensitive C32 cell line, for which <60 cells survived following treatments with the combination of vemurafenib and trametinib) from each of the 102 tested conditions, covering the entire panel of 17 cell lines, 3 drugs (DMSO, vemurafenib alone, and vemurafenib plus trametinib) and 2 timepoints. Log-transformed single-cell data for multiplexed measurements of differentiation state markers, MITF, NGFR and AXL, and for multiplexed measurements of MAPK signaling, p-ERK^T202/Y204^, p-S6^S235/S236^ and Ki-67, were processed by z-scoring (for each protein measurement) across all 6069 cells, followed by principal component analysis (PCA) using the MATLAB built-in function ‘pca’. 80% of the variance in the differentiation state single-cell data and 92.5% of the variance in MAPK single-cell data were captured by the first two principal components (PCs). We thus performed *t*-distributed stochastic neighbor embedding (t-SNE) analysis on the scores from the first two PCs of each of the single-cell datasets by applying the built-in ‘tsne’ function in MATLAB 2019b. We used the ‘barneshut’ algorithm, learning rate of 1000 for the optimization process, a maximum number of optimization iterations of 2000, perplexity of 480, and exaggeration factor of 4. To analyze the baseline state and treatment-induced changes in the single-cell behavior of each cell line, we used the t-SNE map overlaid with the projection of z-scored single-cell measurements (color coded between blue and red) for that cell line while showing data for other cell lines in gray.

To compare baseline and treatment-induced heterogeneities in the expression of each protein, we evaluated Fano factor (a.k.a index of dispersion) by computing the variance-to-mean ratio for single-cell measurements across each population of cells. To compare single-cell heterogeneity in multiple differentiation state markers (MITF, NGFR and AXL) simultaneously, we determined the average cell-to-cell distance by computing the pairwise Euclidean distance among individual cells (after z-scoring the data for each protein marker), and the mean of all possible pairwise distances were reported for each population of cells.

### Measurements of growth rate, drug sensitivity and combination effectiveness

Growth rate inhibition assays were performed in 96-well clear bottom black plates (Corning, Cat# 3904). Cells were counted using a TC20 automated cell counter (Bio-Rad Laboratories) and seeded in 200 µl of full growth media at a density of 1,200-5,000 cells per well, depending on the baseline proliferation rate of each cell line. Using a D300e Digital Dispenser, cells were treated the next day with small molecule compounds at reported doses or vehicle (DMSO). Measurements of live cell count across multiple timepoints were then used to calculate the net growth rate for each treatment condition. To measure the number of surviving cells at each timepoint, cells were fixed in 4% paraformaldehyde for 20 min at room temperature. Cells were washed twice with PBS and incubated with Hoechst 33342 (Thermo Fisher, Cat# H3570, 1:20000) for 20 min at room temperature. Cells were washed again with PBS and imaged with a 10x objective using an ImageXpress Micro Confocal High-Content Imaging System (Molecular Devices) or an Operetta CLS High-Content Imaging System (Perkin Elmer). A total of 9 sites were imaged per well. Nuclear segmentation and cell counting was performed using CellProfiler ^47^.

The net growth rate for cells treated with individual compounds (including MAPK inhibitors or epigenetic inhibitors), or their combination, or vehicle, was calculated from time-dependent changes in live cell count according to the following equation:

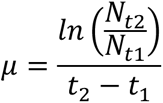

where *N_t1_* and *N_t2_* represent the number of cells measured at timepoints *t* = *t_1_* and *t* = *t_2_*, respectively, and *μ* describes the net growth rate of cells during the time period between *t_1_* and *t_2_*. Average net growth rates for each treatment condition were calculated as the mean of growth rates measured across multiple consecutive timepoints. To compare drug sensitivity among cell lines while correcting for differences in their baseline proliferation rates, drug-induced normalized growth rates (a.k.a. DIP rates ^37^) were computed as follows:

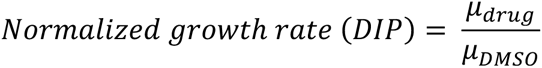

As shown above, normalized growth rate for cells treated with a specific drug is calculated by normalizing the average net growth rate measured for drug-treated cells (*μ_drug_*) to that measured for vehicle (DMSO)-treated cells (*μ_DMSO_*). Normalized growth rates < 0 indicate a net cell loss (i.e. cytotoxicity), a value of 0 represents no change in viable cell number (i.e. cytostasis), a value > 0 indicates a net cell gain (i.e. population growth), and a value of 1 represents cell growth at the same rate as in vehicle-treated (control) cells.

To quantify the benefit resulting from combining two drugs, e.g. an epigenetic inhibitor (A) with a MAPK inhibitor (B), we evaluated their interactions based on the Bliss independence model ^38^. We first calculated the fraction of cells affected by each treatment as follows:

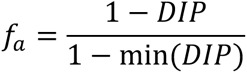

where min (*DIP*) represents the minimum of DIP values reported across all cell lines and drug treatment conditions in this study. Using *f_a_*, we overcome the Bliss metric limitation for the analysis of unbounded drug effects such as the normalized growth rates ^48^, while highlighting combined interactions that influence drug efficacy, a parameter that is affected (more obviously than potency) by cell-to-cell heterogeneity and the presence of small subpopulations of drug-tolerant cells ^49^. We then used measurements of *f_a_* for individual and combination treatments to compute the deviation from Bliss Independence (DBI) as follows:

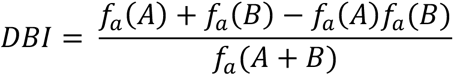

where *f_a_*(*A*) and *f_a_*(*B*) represent the fraction of cells affected by each drug separately, and *f_a_*(*A*+*B*) represents that of the combination treatment (A plus B). Based on this definition, the calculated DBI compares the observed response to that expected given independent action for the two individual treatments. DBI = 1 represents an independent (additive) effect equal to what is expected for the combination of drugs that act independently, DBI < 1 represents a combined effect stronger than expected for an independent combination (i.e. synergism), and DBI > 1 represents a combined effect weaker than expected for an independent combination (i.e. antagonism).

### Epigenetic compound screen

COLO858 and MMACSF cells were counted using a TC20 automated cell counter (Bio-Rad Laboratories) and seeded in 96-well clear bottom black plates at 2000 and 3000 cells per well (excluding 36 wells at the edges), respectively. Using the HP D300e Digital Dispenser, cells were treated the next day with either vehicle (DMSO or water) or two different doses (0.2 µM and 1 µM) of each of the 276 compounds in an epigenetic compound library. After 24 h, cells were either fixed or treated with vemurafenib alone (at 100 nM), or vemurafenib (at 100 nM) plus trametinib (at 10 nM), or vehicle (DMSO). Cells were then grown for a further 72 h or 120 h prior to fixation. All experimental conditions with an epigenetic compound treatment (at each dose and timepoint) were tested in two replicates. Experimental conditions that did not include an epigenetic compound treatment (i.e. treatments with only vehicle, vemurafenib, or vemurafenib in combination with trametinib) were repeated in all 96-well plates during the entire period of compound screening, creating a total of 276 replicates that were used to evaluate plate-to-plate and day-to-day experimental robustness. To differentiate the impact of epigenetic compounds on each cell line at different states of MAPK signaling, the growth rates for cells treated with each dose of an epigenetic compound were compared with growth rates for cells treated without any epigenetic compound, following 72 h and 120 h of exposure to each of the MAPK inhibitor treatment conditions (i.e., DMSO, vemurafenib, and vemurafenib plus trametinib). Significant epigenetic compounds were defined as those that led to a statistically significant decrease in normalized growth rate (with an effect size of at least 0.5 day^-1^) in at least one of the tested conditions (i.e. cell lines, timepoints, or MAPK inhibitor conditions). Statistical significance was evaluated as *P* < 0.05 based on two-sample *t*-test following correction for multiple comparisons using the Dunn-Sidak method. Additional details about the epigenetic compound screen are available in Supplementary Table 2.

### Hierarchical clustering

To infer potential variations in the mechanisms of action of the selected group of 58 epigenetic compounds, treatment-induced changes in growth rate, p-ERK^T202/Y204^, p-Rb^S807/811^, and MITF across different cell lines and MAPK inhibitor conditions were integrated into a matrix for unsupervised clustering. Growth rates for epigenetic inhibitor conditions were averaged across three timepoints and two doses of the inhibitor and their differences relative to the time-averaged growth rate in the absence of any epigenetic compound were used for each MAPK inhibitor treatment condition (i.e., DMSO, vemurafenib, and vemurafenib plus trametinib). For p-ERK^T202/Y204^, p-Rb^S807/811^, and MITF, log-transformed signal intensities (from quantitative immunofluorescence microscopy) were averaged across three timepoints and two doses of the inhibitor and their differences relative to the corresponding MAPK inhibitor condition in the absence of any epigenetic compound were used. Unsupervised hierarchical clustering of the integrated data was then carried out using MATLAB 2018b with the Correlation distance metric and the Complete (farthest distance) algorithm for computing the distance between clusters.

### Statistical analysis of drug response data

All data with error bars were presented as mean values ± standard deviation (s.d.) using indicated numbers of replicates. The significance of pairwise correlations among drug response data were evaluated based on *P* values associated with the corresponding Pearson’s correlation coefficients (*r*). The statistical significance of the effects of epigenetic compounds (in the first stage of screen) was evaluated based on *P* < 0.05 generated from two-sample *t*-test following correction for multiple comparisons using the Dunn-Sidak method. To identify the statistical significance of differences between mean or median of measurements within two different groups, *P* values from two-sample *t* test or from nonparametric Mann-Whitney U-test were used, respectively. Statistical significance of the difference in % change in tumor size in mouse experiments was determined by two-way analysis of variance (ANOVA). Statistical analyses were performed using MATLAB 2019b software.

### Partial least square regression (PLSR) analysis

We used partial least square regression (PLSR) analysis ^17,40,50^ to generate models that linked epigenetic treatment-induced changes in normalized growth rates to baseline and treatment-induced changes in signaling and phenotypic data. By combining data for each epigenetic inhibitor across eight cell lines, we generated one model to predict responses to each epigenetic inhibitor, when used either individually or in combination with BRAF/MEK inhibitors. “Response” variables for each model were defined as normalized growth rates (averaged across 3-5 days of treatment) in the absence or presence of that compound (together with either DMSO, vemurafenib alone, or vemurafenib in combination with trametinib) across the eight cell lines. “Input” vectors were constructed by combining measurements of a total of 8 proteins (including signaling proteins, differentiation state and phenotypic markers) at the baseline (i.e. either DMSO, vemurafenib alone, or vemurafenib in combination with trametinib), following epigenetic inhibitor treatment (together with either DMSO, vemurafenib alone, or vemurafenib in combination with trametinib), and their changes induced by the epigenetic inhibitor which were computed by taking the ratio of the two sets of measurements. All protein measurements were performed by quantitative immunofluorescence microscopy, log-transformed and averaged cross three timepoints (including 24 h after epigenetic inhibitor or DMSO treatment, followed by 72 h and 120 h of MAPK inhibitor or DMSO treatment). The data were then z-scored across all conditions and cell lines prior to the application of PLSR analysis using the built-in MATLAB function ‘plsregress’.

To evaluate the predictability of the linear relationship between the input and output variables in each model, we used leave-one-out cross validation. The goodness of fit for each model was calculated using R^2^. Prediction accuracy was evaluated by Q^2^ and the *P* values generated from pairwise Pearson’s correlations between the measured and predicted responses following cross-validation. For the assessment of relative variable importance in each PLSR model, the information content of each variable was assessed by its variable importance in the projection (VIP) ^41^.

### Multi-linear regression (MLR) analysis

To statistically test the hypothesis that the relative levels of KDM4B, ZNF217 and differentiation state markers NGFR and AXL can *predict* the selective sensitivity of melanoma cells to JIB-04 and SP2509, we performed multi-linear regression (MLR) analysis using the entire panel of 21 cell lines, which were divided into a test set of 14 (to tarin models) and a validation set of 7 cell lines (to evaluate the model performance). To group the cell lines into training and validation sets, we first sorted the 21 cell lines based on their sensitivity to each compound. Beginning from the most sensitive or resistant cell line, we picked 2 cell lines for the training set and 1 cell line for the validation set, then another 2 cell lines for the training set and 1 cell line for the validation set, etc. We then used the training set to develop MLR models of treatment efficacy for SP2509 and JIB-04 using protein levels of NGFR, AXL, KDM4B and ZNF217 measured by quantitative immunofluorescence microscopy. “Response” variables were defined as normalized growth rates (averaged across 3-5 days of treatment) in the presence of either SP2509 (at 1 µM) or the combination of JIB-04 (at 0.2 µM), vemurafenib (at 100 nM) and trametinib (at 10 nM). “Input” vectors were constructed by combining baseline (drug-naïve) measurements of the fractions of NGFR^High^, AXL^High^, KDM4B^High^ and ZNF217^High^ cells in each cell line. We then used the built-in MATLAB function ‘regress’ to return a vector of coefficient estimates for an MLR of the responses to each treatment condition. These coefficients were then used to predict the responses of the validation cell lines to each treatment using an input vector generated for those cell lines. To evaluate the predictability of the models, we evaluated the relationship between the measured and predicted responses for the validation set of cell lines using Pearson’s correlation analysis. The goodness of fit for each model was calculated using R^2^ of measured versus fitted responses for models generated using all 21 cell lines.

### Gene knockdown by siRNA

In the first round of siRNA-mediated knockdown experiments, WM115, WM902B and A2058 cells were seeded in 200 µl of antibiotic-free growth media in 96-well clear bottom black plates at seeding densities of 3500, 4000, and 1700 cells per well, respectively. After 24 hours, cells were transfected using the DharmaFECT 1 transfection reagent (GE Dharmacon T-2001-01) with Dharmacon’s ON-TARGETplus Human SMARTpool siRNAs, including four target-specific siRNAs combined into a single pool to increase the likelihood of effective gene silencing. The SMARTpool siRNAs included KDM4A (L-004292-00-0005), KDM4B (L-004290-00-0005), KDM5A (L-003297-02-0005), KDM5B (L-009899-00-0005), or non-targeting control (D-001810-10-05) and were used at 25 nM. To rule out the possibility of off-targeting by KDM4B siRNAs, we then examined separately the effects of three constituent ON-TARGETplus KDM4B siRNAs (J-004290-08-0005, J-004290-09-0005, J-004290-11-0005) at 25 nM. To evaluate the effect of BRAF/MEK inhibition in cells in which either of the proteins were knocked down, WM115 and WM902B cells were further treated with vemurafenib (at 100 nM) plus trametinib (at 10nM) or DMSO 48 h following siRNA addition. After another 48 h, cells were fixed and analyzed using quantitative immunofluorescence microscopy.

### Gene knockout by CRISPR-Cas9

A2058 cells were seeded in a 24-well plate at a seeding density of 30,000 cells per well for 24 hours. The next day, cells were treated at MOI of 0.3 with the pre-designed Edit-R lentiviral Cas9 nuclease (Dharmacon, Cat# VCAS10124). Polybrene (at 8 μg/ml) was added to enhance efficiency of the viral infection. Following an incubation period of 72 h with the lentivirus, Cas9-lentiviral-treated cells were selected with blasticidin over a period of 1.5 weeks. Surviving A2058 Cas9-positive cells were subsequently infected with either non-targeting control (Dharmacon, Cat# VSGC10216) or three independent KDM1A lentiviral sgRNAs (Dharmacon, Cat# VSGH10142-246522030, VSGH10142-246999191, and VSGH10142-246522028). The efficiency of KDM1A knockout and the consequential changes in cell viability were measured via immunostaining for the KDM1A protein at 96 h, followed by quantitative immunofluorescence microscopy.

### *In vivo* xenograft assays

All mouse experiments were carried out in accordance with procedures approved by the Institutional Animal Care and Use Committee (IACUC) at the University of Michigan. Athymic, 5-6 weeks old female nude (NU/J) mice were purchased from The Jackson Laboratory. Mice were maintained and housed under standard animal housing protocol at the University of Michigan Unit for Laboratory Animal Care (ULAM). For xenograft tumor injections, mice were first anesthetized using 3% vaporized isoflurane. 2.5 × 10^6^ cells from either of the melanoma cell lines A375 or WM2664 suspended in 200 μl of growth factor-reduced Matrigel (Thermo Fisher, CB-40230C) in PBS (1:1) were injected subcutaneously in the right flank of each mouse. Tumor xenografts were monitored three times a week with digital calipers. Once the tumors were palpable and the mean volume across all tumors reached a volume of approximately 170 mm^3^, the mice were randomly allocated to two treatment groups of 5 mice per group. SP2577 (at a daily dose of 80 mg/kg) or vehicle (20% DMSO, 20% Cremophor EL, plus 60% sterile water) was then administered to each mouse via I.P. injection of a solution of 200 μl. Mouse body weight, tumor volume, and general health were monitored three times a week for 12 days. At the endpoint of the study, mice were euthanized by CO_2_ asphyxiation and bilateral pneumothorax.

### Data availability

Source data for all high-throughput experiments, including growth rate inhibition assays, epigenetic compound screening, single-cell profiling of MAPK signaling and differentiation state in melanoma cells, as well as profiling of signaling and phenotypic responses to epigenetic inhibitors are presented in Supplementary Datasets 1-14. Supplementary Dataset 1 is associated with data presented in Fig. 1a, b. Supplementary Dataset 2 is associated with data presented in Fig. 1c, d. Supplementary Dataset 3 is associated with data presented in Fig. 1e, f. Supplementary Dataset 4 is associated with data presented in Fig. 2b. Supplementary Dataset 5 is associated with data presented in Fig. 2c. Supplementary Dataset 6 is associated with data presented in Fig. 3 and 4. Supplementary Datasets 7-10 are associated with data presented in Fig. 5a. Supplementary Datasets 11-14 are associated with data presented in Fig. 6d.

### Code availability

Programming was performed in MTALAB using built-in functions and parameters as described in detail is Methods. Custom analysis MATLAB scripts are available upon request.

## Acknowledgments

We thank Meenhard Herlyn for advice. We also thank Noah Brown, LaJuanda Carter and members of the Fallahi-Sichani laboratory for help and discussion. This work was supported by the University of Michigan, University of Virginia, and awards from the Elsa Pardee Foundation and V Foundation for Cancer Research V2017-011, Department of Defense PRCRP Career Development Award W81XWH1810427, NIH grants R00-CA194163 and R35-GM133404 (to MFS), P30-CA046592 (University of Michigan Rogel Cancer Center), P30-CA044579 (University of Virginia Cancer Center), NCI Training Grant award T32-CA009676 (to MK), the Welch Foundation I-1878 and CPRIT RP160493 (to EDM) and NCI SPORE grant P50-CA070907.

## Author Contributions

M.K. designed and performed the experiments, analyzed the experimental data, performed analyses, and wrote the manuscript. M.M. designed and performed the experiments. E.D.M. synthesized reagents for the experiments. M.F.-S. designed the experiments, performed analyses and wrote the manuscript.

## Extended Data Figure Legends

**Extended Data Figure 1.**
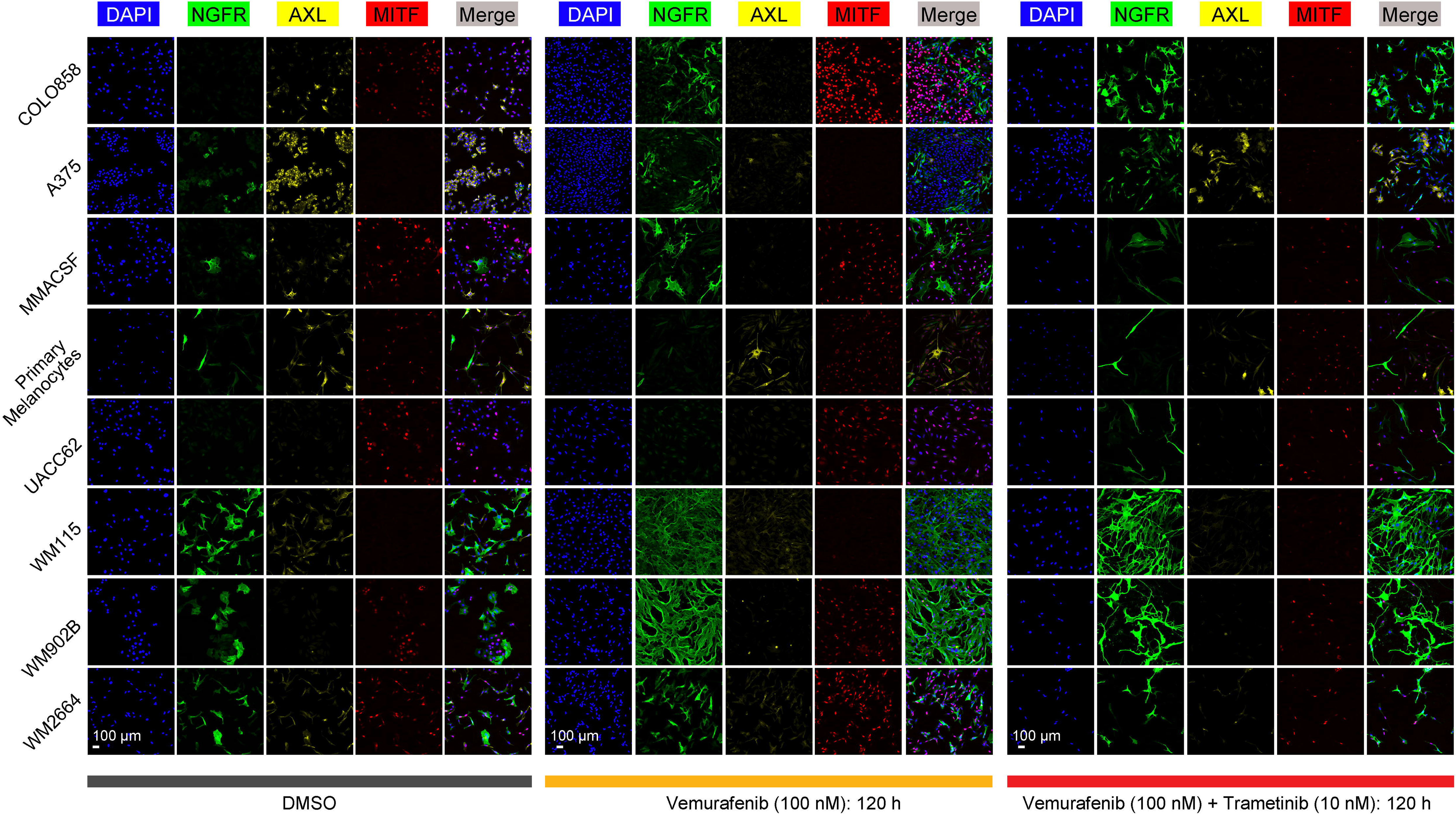
Representative multiplexed immunofluorescence images of differentiation state markers MITF, NGFR and AXL, co-stained in indicated melanoma cell lines or primary melanocytes before and after treatment with BRAF/MEK inhibitors at indicated doses and timepoints. Scale bars represent 100 μm.

**Extended Data Figure 2.**
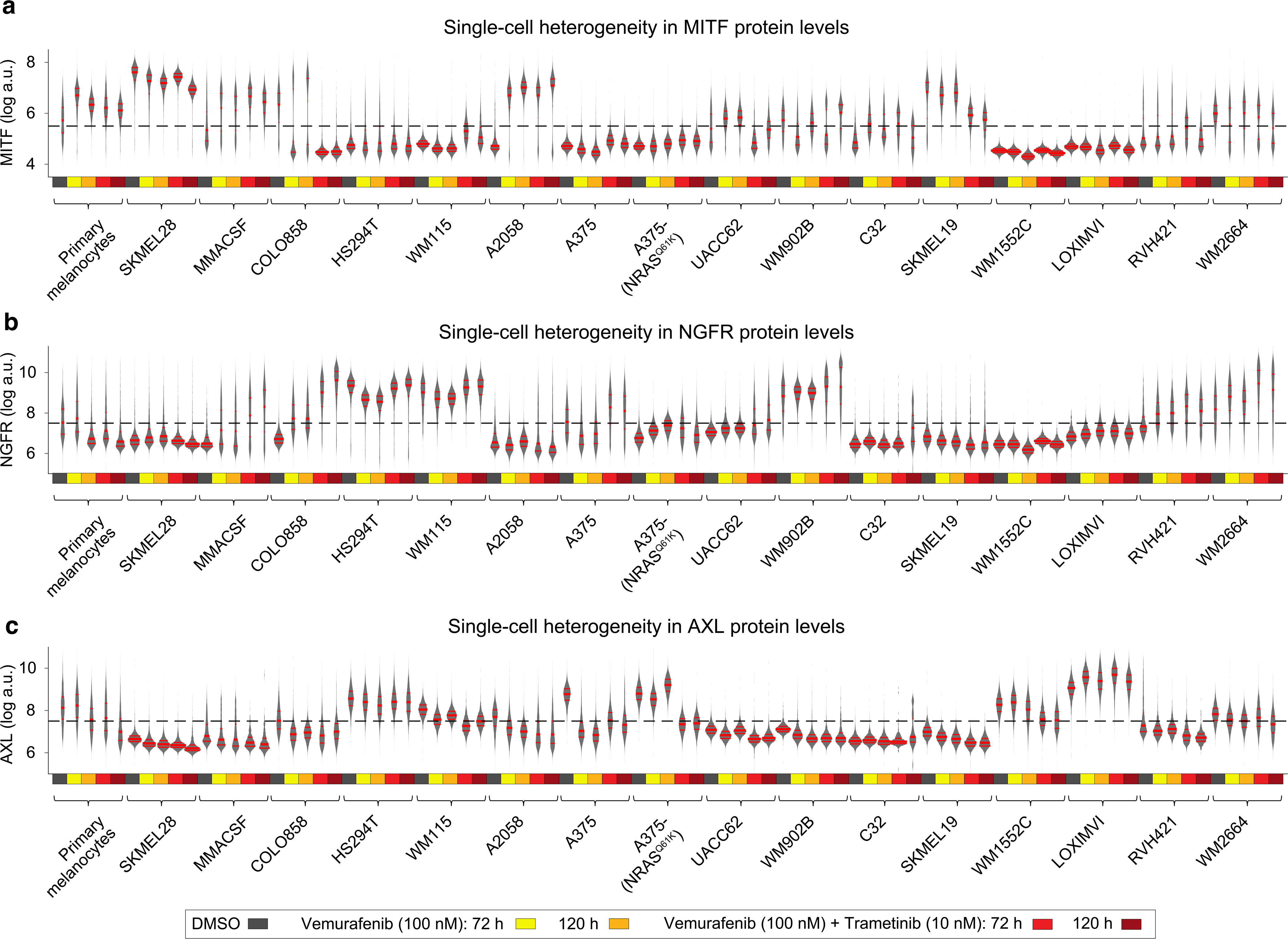
Single-cell heterogeneity in differentiation state markers MITF, NGFR and AXL, quantified by multiplexed immunofluorescence imaging across 16 melanoma cell lines and primary melanocytes before and after treatment with BRAF/MEK inhibitors at indicated doses and timepoints. The distributions of single-cell data across different conditions are shown by violin plots, highlighting the median and interquartile (25% and 75%) ranges.

**Extended Data Figure 3.**
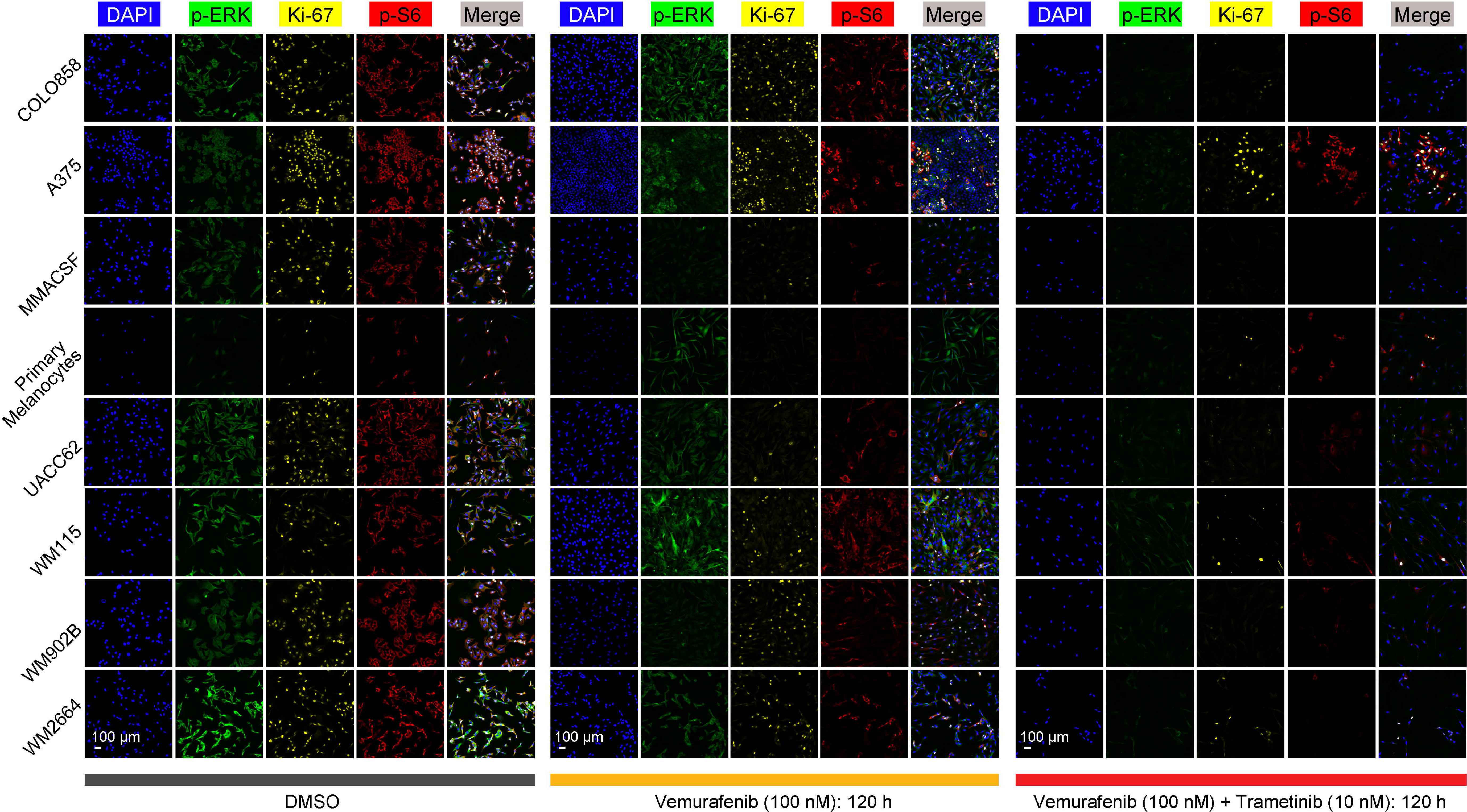
Representative multiplexed immunofluorescence images of p-ERK^T202/Y204^, p-S6^S235/S236^ and Ki-67 proteins, co-stained in indicated melanoma cell lines or primary melanocytes before and after treatment with BRAF/MEK inhibitors at indicated doses and timepoints. Scale bars represent 100 μm.

**Extended Data Figure 4.**
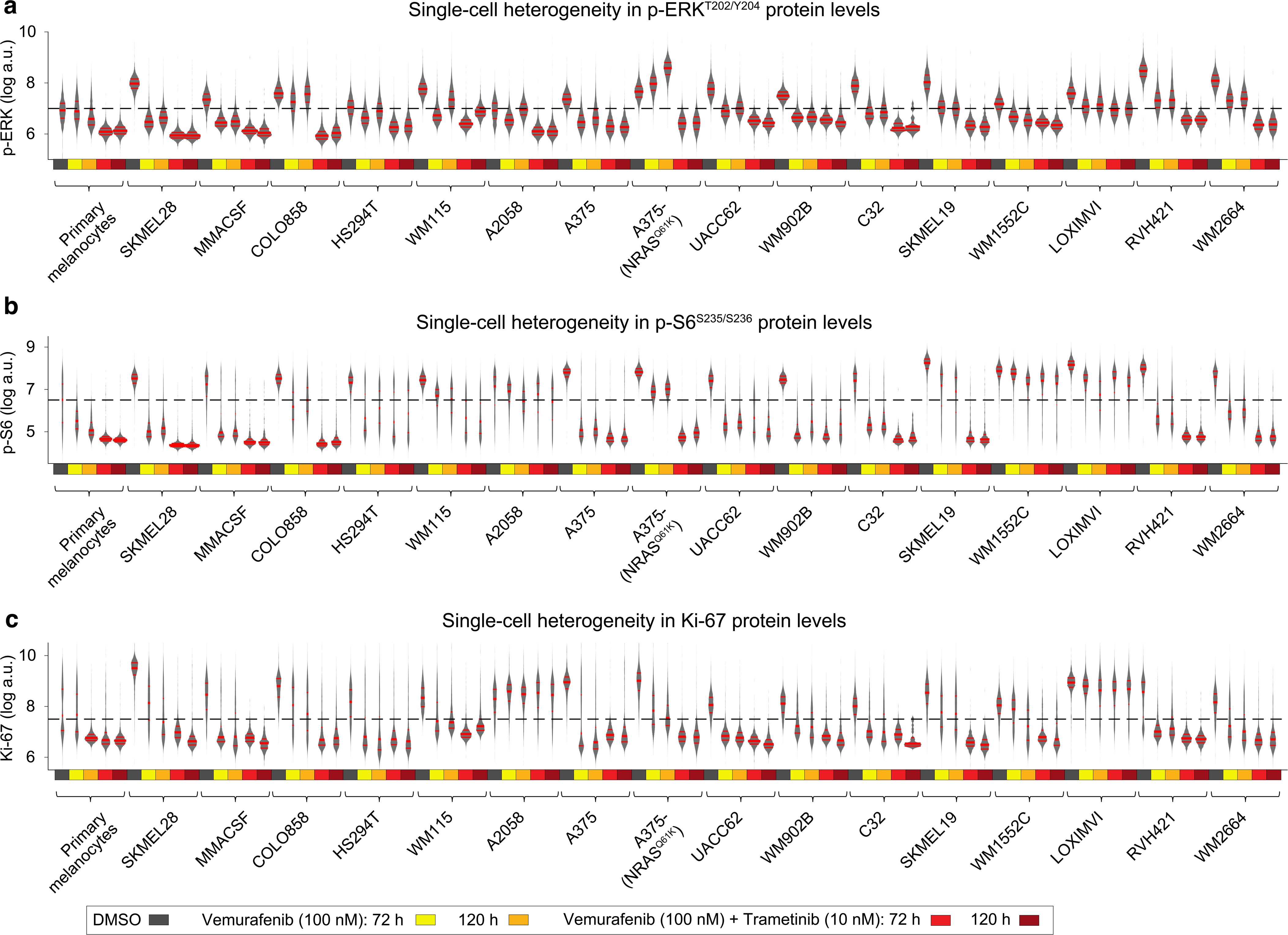
Single-cell heterogeneity in p-ERK^T202/Y204^, p-S6^S235/S236^, and Ki-67 proteins, quantified by multiplexed immunofluorescence imaging across 16 melanoma cell lines and primary melanocytes before and after treatment with BRAF/MEK inhibitors at indicated doses and timepoints. The distributions of single-cell data across different conditions are shown by violin plots, highlighting the median and interquartile (25% and 75%) ranges.

**Extended Data Figure 5.**
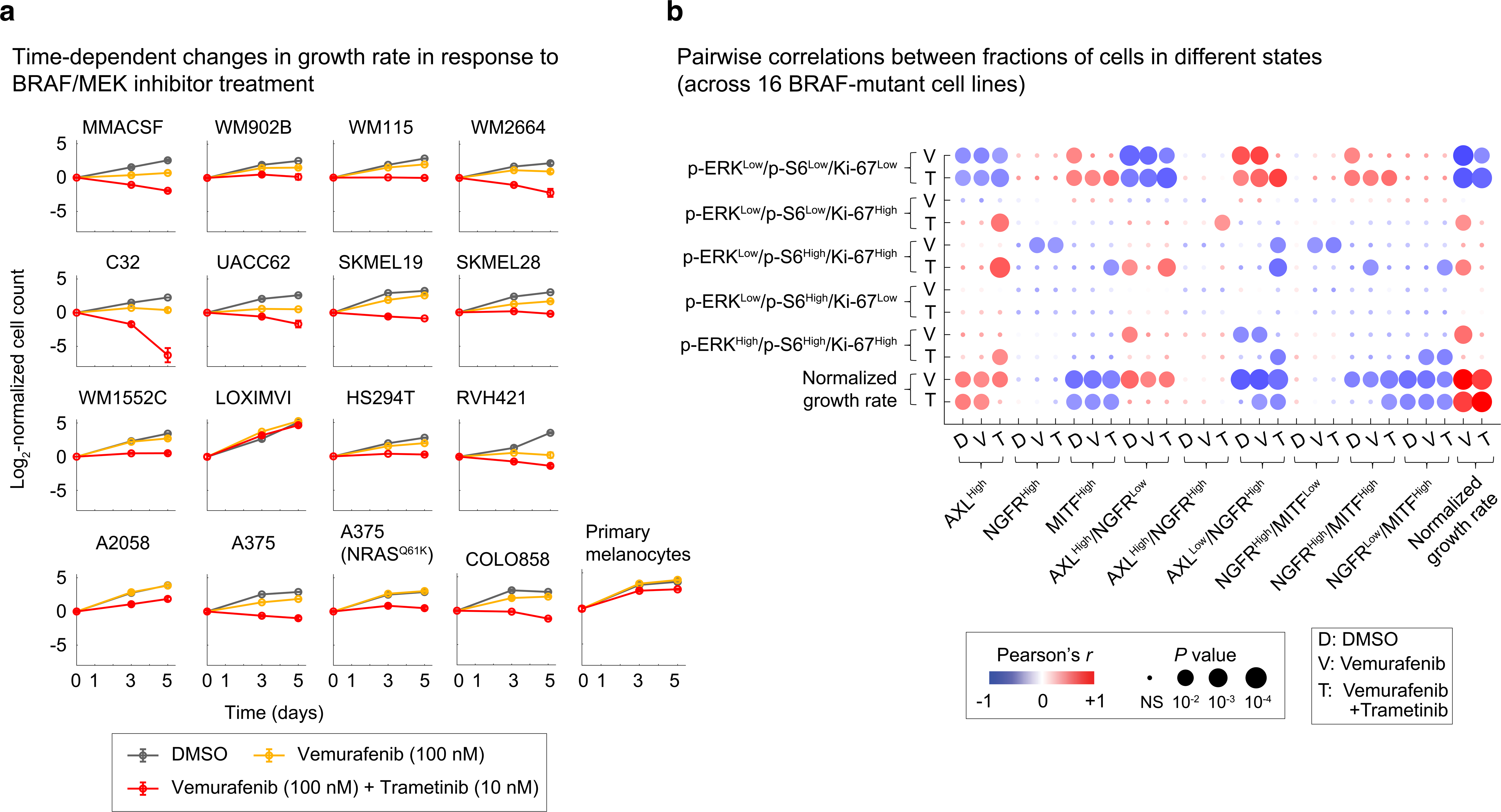
Single-cell analysis uncovers heterogeneities in melanoma differentiation, proliferation and MAPK signaling states across a wide range of BRAF/MEK inhibitor sensitivity. (a) Log_2_-normalized changes in live cell count across three timepoints (including 0, 3, and 5 days) following exposure of 16 BRAF^V600E/D^ melanoma cell lines and primary melanocytes to DMSO, vemurafenib (at 100 nM) or vemurafenib (at 100 nM) plus trametinib (at 10 nM). Data represent mean values ± s.d. calculated from two replicates per treatment condition. (b) Pairwise Pearson correlation analysis between diverse differentiation states, MAPK signaling states, and normalized growth rates cross 16 melanoma cell lines treated with BRAF inhibitor (vemurafenib at 100 nM) or the combination of BRAF and MEK inhibitors (vemurafenib at 100 nM and trametinib at 10 nM) for 3-5 days.

**Extended Data Figure 6.**
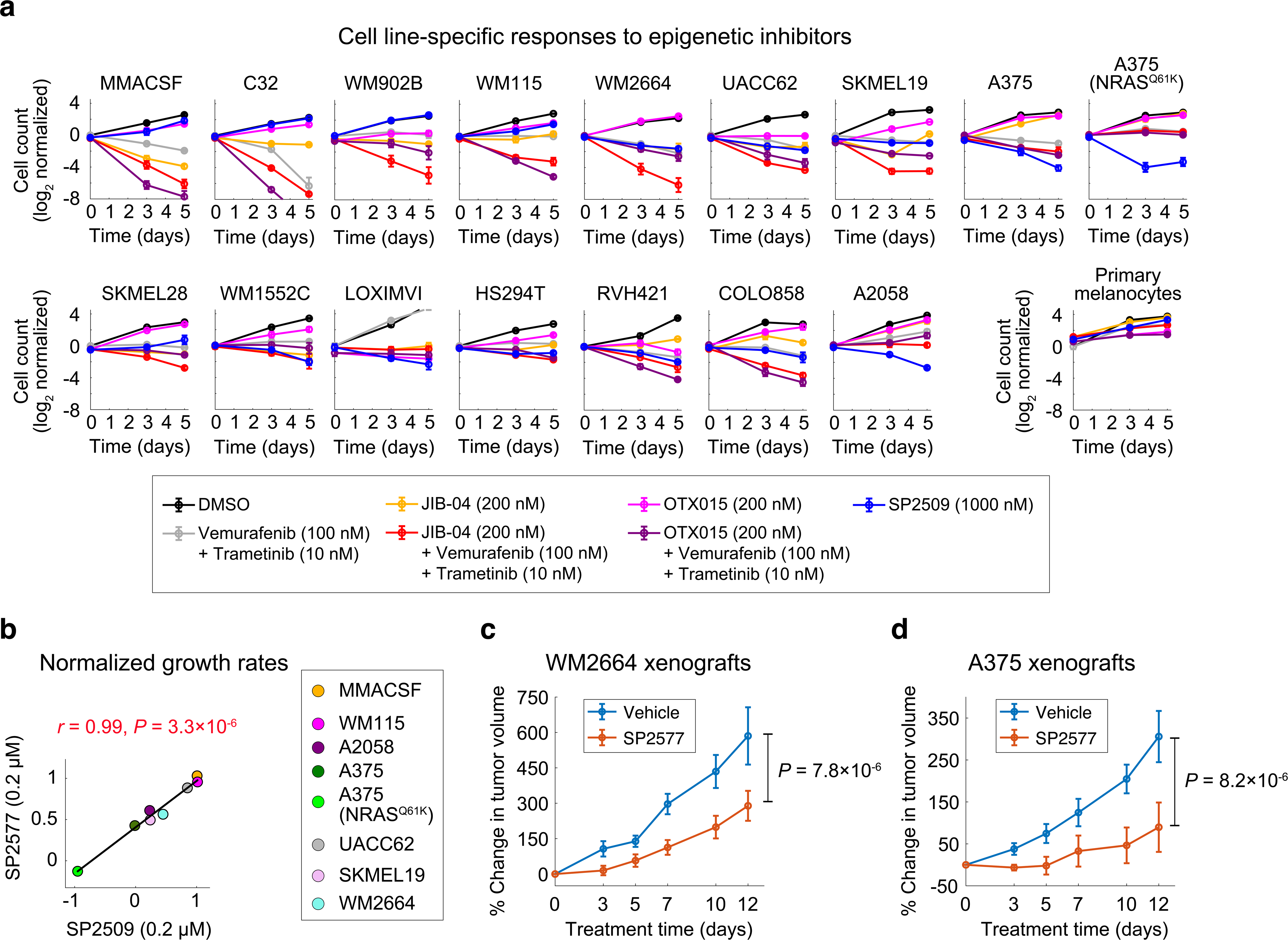
Comparing treatment-induced changes in growth rates of melanoma cells following pharmacological inhibition of KDM1A, Jmj-KDMs and BET proteins. (a) Log_2_- normalized changes in live cell count following exposure of melanoma cell lines and non-transformed primary melanocytes to different drugs at indicated doses for a period of 5 days. Data represent mean values ± s.d. calculated from two replicates per treatment condition. (b) Pairwise Pearson correlation between the effect (i.e. normalized growth rates) of KDM1A inhibitor SP2509 (at 0.2 μM) and its clinical formulation, SP2577 (at 0.2 μM) across eight BRAF-mutant melanoma cell lines. (c,d) Mice bearing xenografts of WM2664 cells (c) and A375 cells (d) were treated as shown by the KDM1A inhibitor SP2577 or vehicle for a period of 12 days, to determine the effect on tumor growth. Data represent mean values ± s.e.m across 5 mice per group. Statistical significance was determined by two-way ANOVA.

**Extended Data Figure 7.**
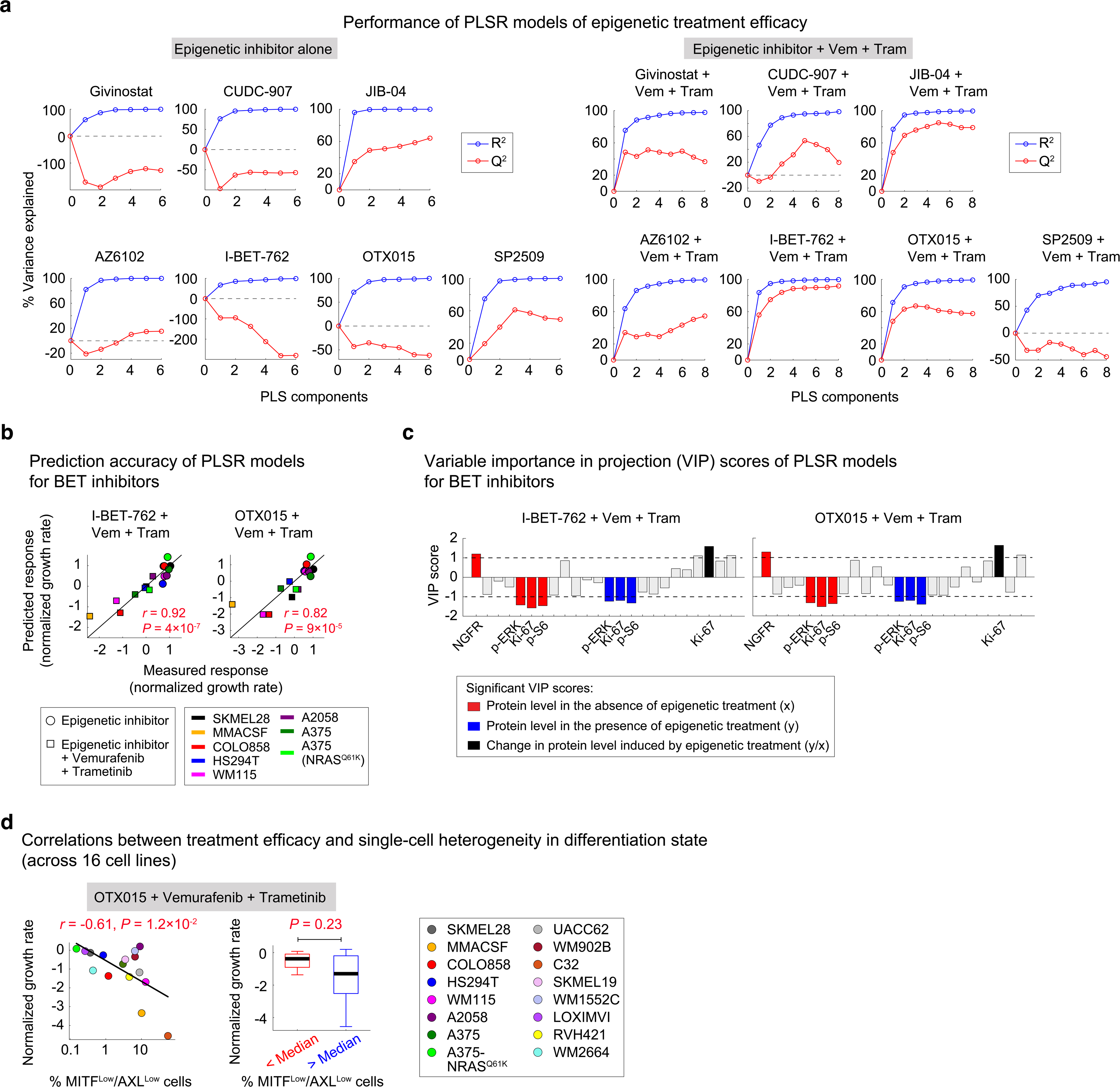
Multivariate modeling identifies differentiation state-specific predictors of epigenetic inhibitor efficacy. (a) Partial least square regression (PLSR) model calibration for the effect of each of the seven epigenetic inhibitors, used individually (left panels) or in combination with vemurafenib and trametinib (right panels), across eight BRAF-mutant melanoma cell lines. R^2^ values (representing goodness of fit) and Q^2^ values (representing prediction accuracy based on leave-one-out cross-validation) are shown for models built with increasing numbers of PLS components. (b) Pairwise Pearson correlation between responses (normalized growth rates) to I-BET-762 (left) or OTX015 (right) in combination with vemurafenib and trametinib measured for each of the eight melanoma cell lines (x-axis) and corresponding responses predicted by PLSR modeling following leave-one-out cross validation (y-axis). (c) PLSR-derived variable importance in the projection (VIP) scores highlighting combinations of protein measurements at the baseline (shown in red), following epigenetic inhibitor treatment (shown in blue), or the ratio of change induced by each epigenetic compound (shown in black), that are predictive of efficacy for I-BET-762 or OTX015 in combination with vemurafenib and trametinib. The sign of VIP score shows whether the change in variable correlated negatively or positively with treatment-induced response. Only VIP scores of greater than 1 or smaller than -1 with a statistically significant Pearson correlation (*P* < 0.05) are highlighted. (d) Association of epigenetic compound efficacy with baseline melanoma differentiation state characterized at a single-cell level across sixteen BRAF-mutant cell lines. The significance of differences between normalized growth rate in response to OTX015 combined with vemurafenib and trametinib versus the baseline fraction of MITF^Low^/AXL^Low^ cells was evaluated by *P* values based on Pearson correlation analysis (left) and nonparametric Mann-Whitney U-test (right). Sixteen cell lines were divided into two groups of eight based on whether the measured variable (% MITF^Low^/AXL^Low^ cells) had a value above or below the median. Data for each group was then presented using box-and-whisker plots and median parameter values and interquartile ranges; bars extending to 1.5× the interquartile range are shown for each condition as a measure of variance.

**Extended Data Figure 8.**
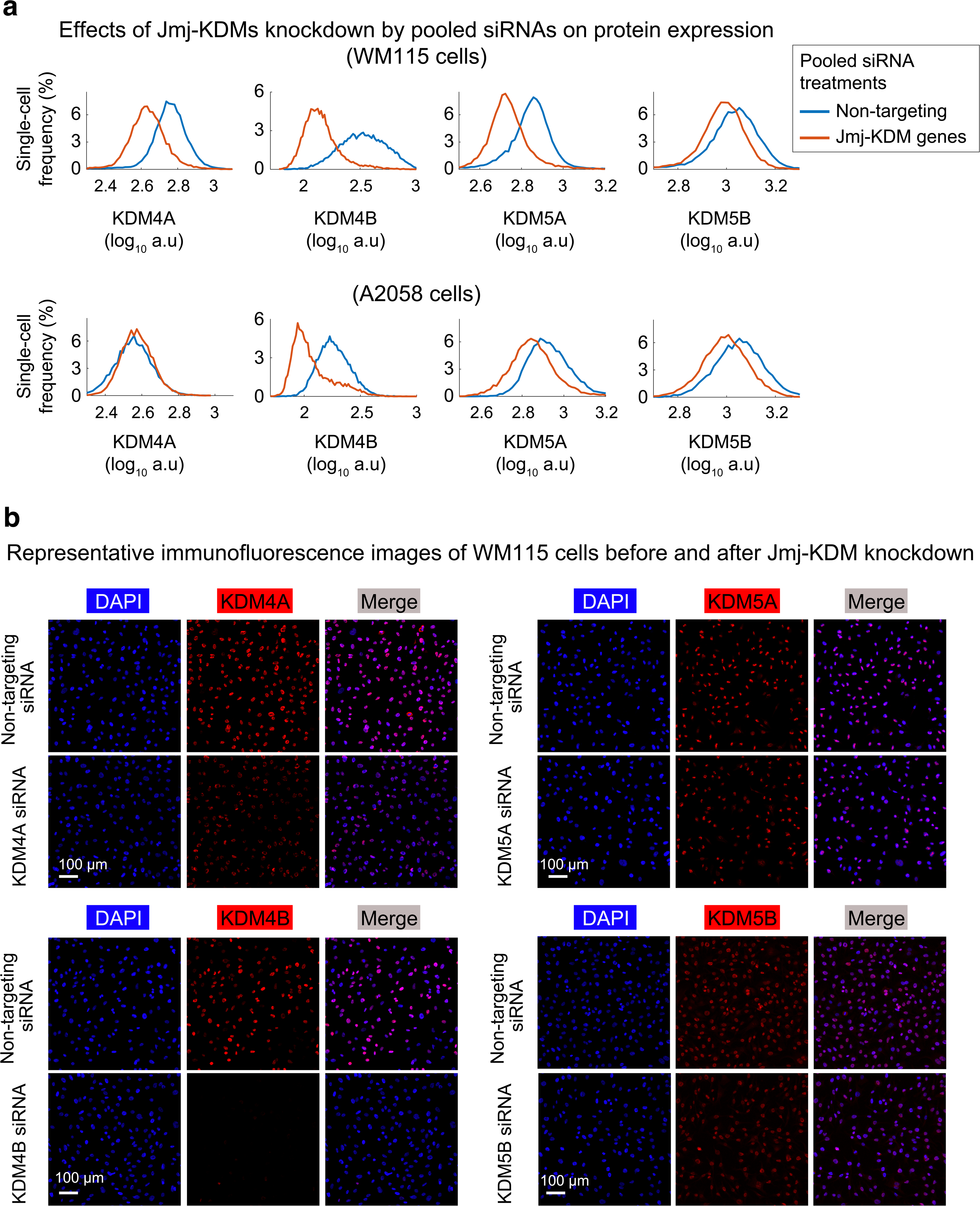
Effects of Jmj-KDM knockdown on the corresponding protein expression levels at a single-cell level. (a) Single-cell protein levels of Jmj-KDM proteins KDM4A, KDM4B, KDM5A, and KDM5B, measured by immunofluorescence microscopy, in WM115 (top) and A2058 cells (bottom) following treatment with pooled siRNAs targeting either KDM4A, KDM4B, KDM5A, or KDM5B, or with non-targeting (control) siRNA for 96 h. (b) Representative immunofluorescence images of WM115 cells before and after each Jmj-KDM protein knockdown. Scale bars represent 100 μm.

**Extended Data Figure 9.**
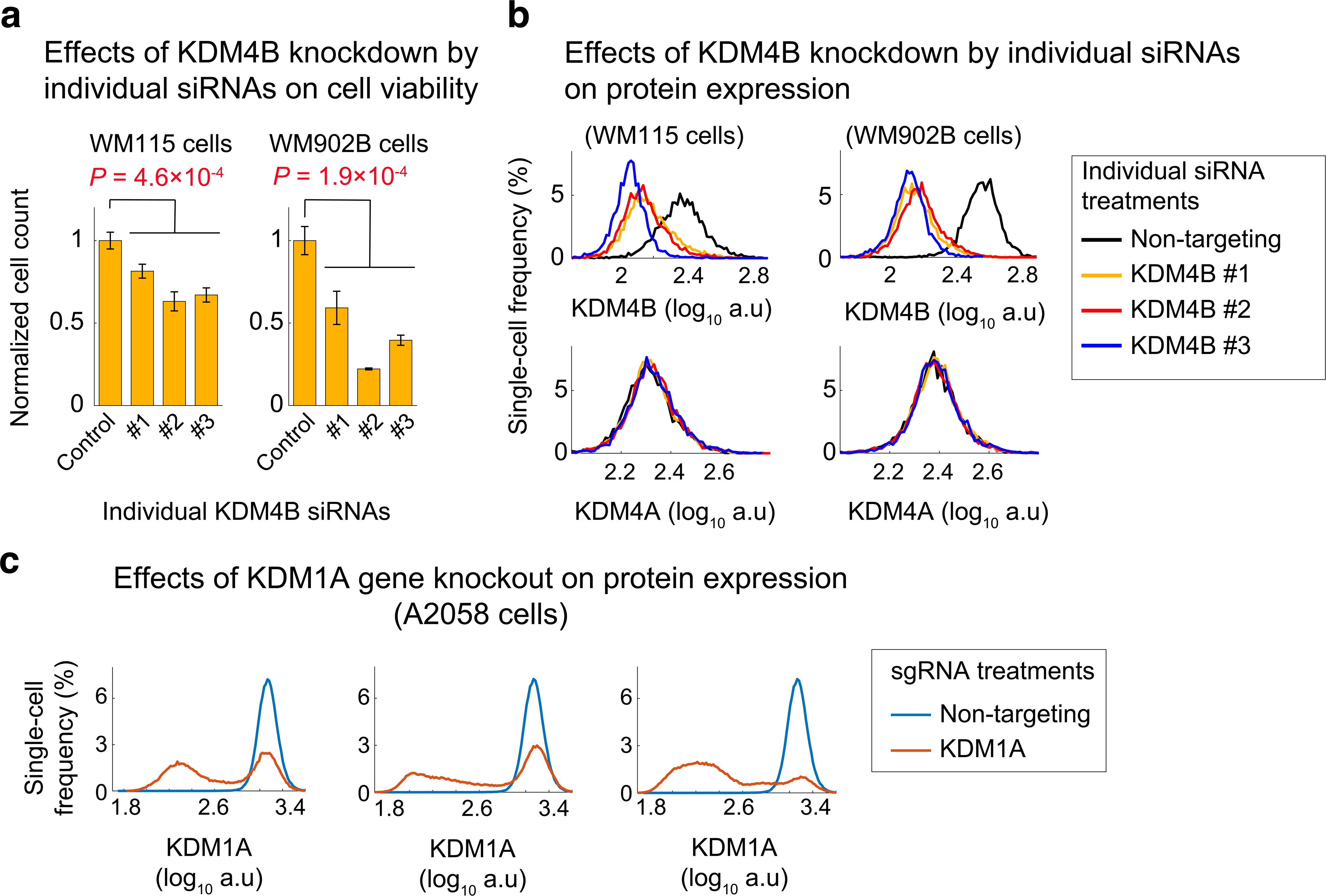
Effects of KDM4B knockdown (by independent siRNAs) and KDM1A CRISPR knockout (by independent sgRNAs). (a) Relative cell viability in WM115 cells (left) and WM902B cells (right) following treatment with three independent siRNAs targeting KDM4B for 96 h. Viability data for each treatment condition were normalized to cells treated with non-targeting (control) siRNA. Data represent mean values ± s.d. calculated across 3 replicates. Statistical significance was determined by two-sided *t* test. (b) Single-cell protein levels of KDM4B (top) and KDM4A (bottom), measured by immunofluorescence microscopy, in WM115 cells (left) and WM902B cells (right) following treatment with three independent siRNAs targeting KDM4B or with non-targeting (control) siRNA for 96 h. (c) Single-cell protein levels of KDM1A, measured by immunofluorescence microscopy, in Cas9-positive A2058 cells following treatment with three different types of KDM1A lentiviral single guide RNA (sgRNA), or with non-targeting (control) sgRNA for 96 h.

**Extended Data Figure 10.**
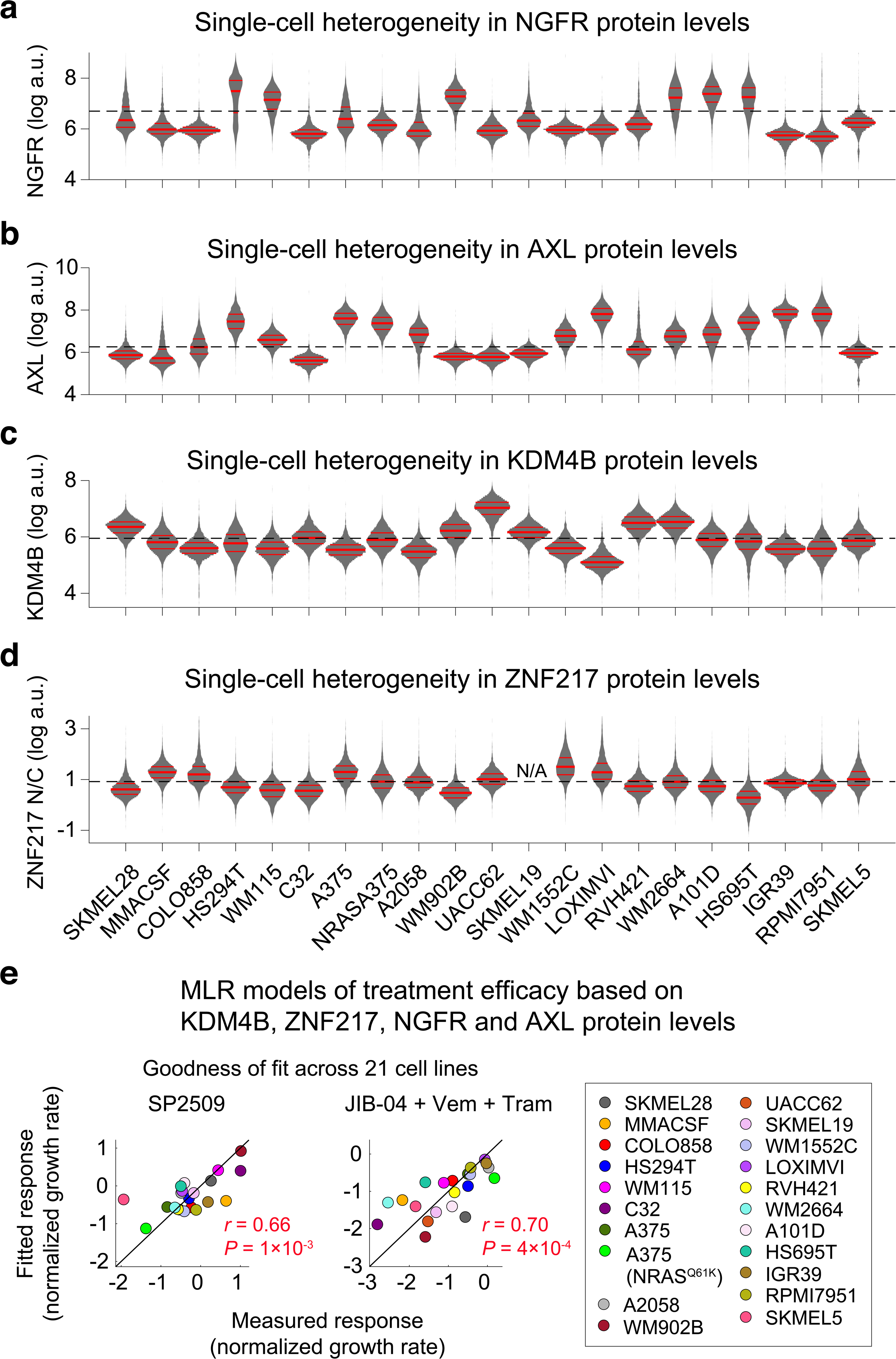
KDM4B and ZNF217 protein levels predict differentiation state-specific sensitivity to JIB-04 and SP2509. (a-d) Single-cell heterogeneity in protein levels of NGFR (a), AXL (b), KDM4B (c) and ZNF217 (d), quantified by multiplexed immunofluorescence imaging across 21 melanoma cell lines. The distributions of single-cell data across different conditions are shown by violin plots, highlighting the median and interquartile (25% and 75%) ranges. (e) Pairwise Pearson correlation between responses (normalized growth rates) to SP2509 (left) or JIB-04 in combination with vemurafenib and trametinib (right), measured for each of the 21 melanoma cell lines (x-axis) and corresponding responses fitted by multi-linear regression (MLR) analysis of all cell lines.

