## Supplementary Information for "Epigenetic modulation reveals differentiation state specificity of oncogene addiction"

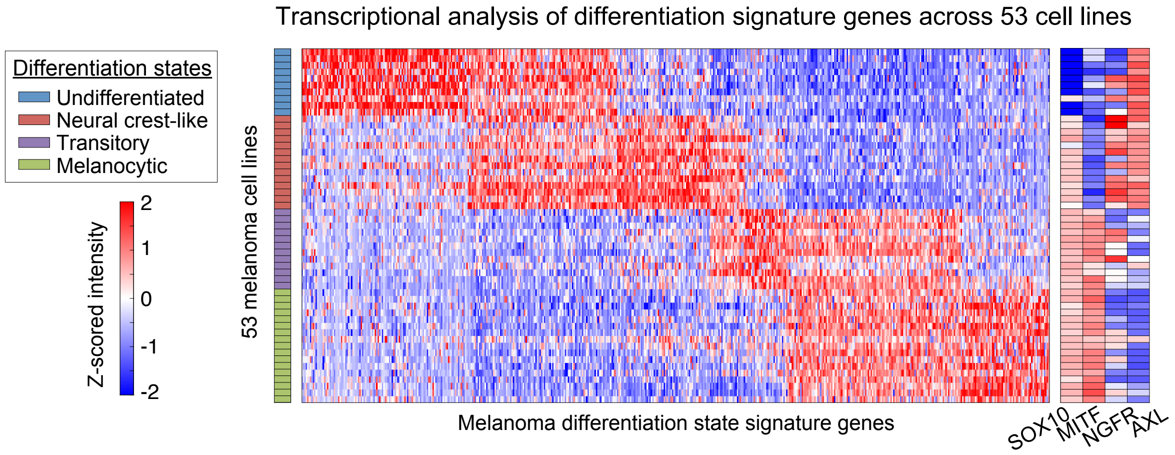


**Supplementary Figure 1. Four melanoma differentiation subtypes revealed by transcriptomic analysis of 53 patient-derived melanoma cell lines from Tsoi *et al*, Cancer Cell 33, 2018.** These states can be distinguished by differentiation state markers SOX10, MITF, NGFR and AXL.

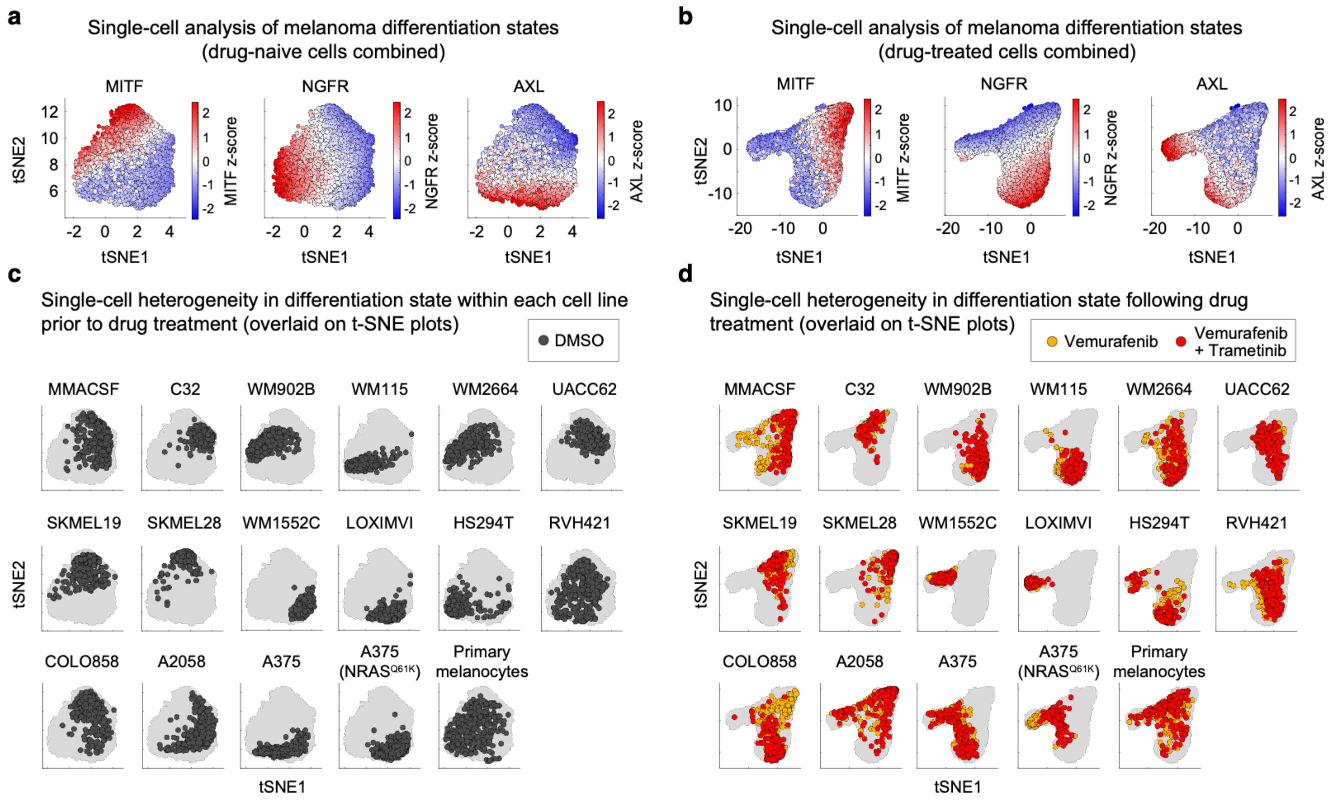


**Supplementary Figure 2. Single-cell analysis uncovers heterogeneities in melanoma differentiation, proliferation and MAPK signaling states. (a,b)** Single-cell protein levels of three melanoma differentiation state markers, MITF, NGFR and AXL, measured by multiplexed immunofluorescence microscopy and visualized by t-SNE, performed separately for drug-naïve cells **(a)** and BRAF/MEK inhibitor-treated cells **(b)**. Experimental conditions and cells utilized for this analysis are the same as those depicted in Figure 1. **(c,d)** Projections of single-cell variations within each individual cell line on t-SNE maps.

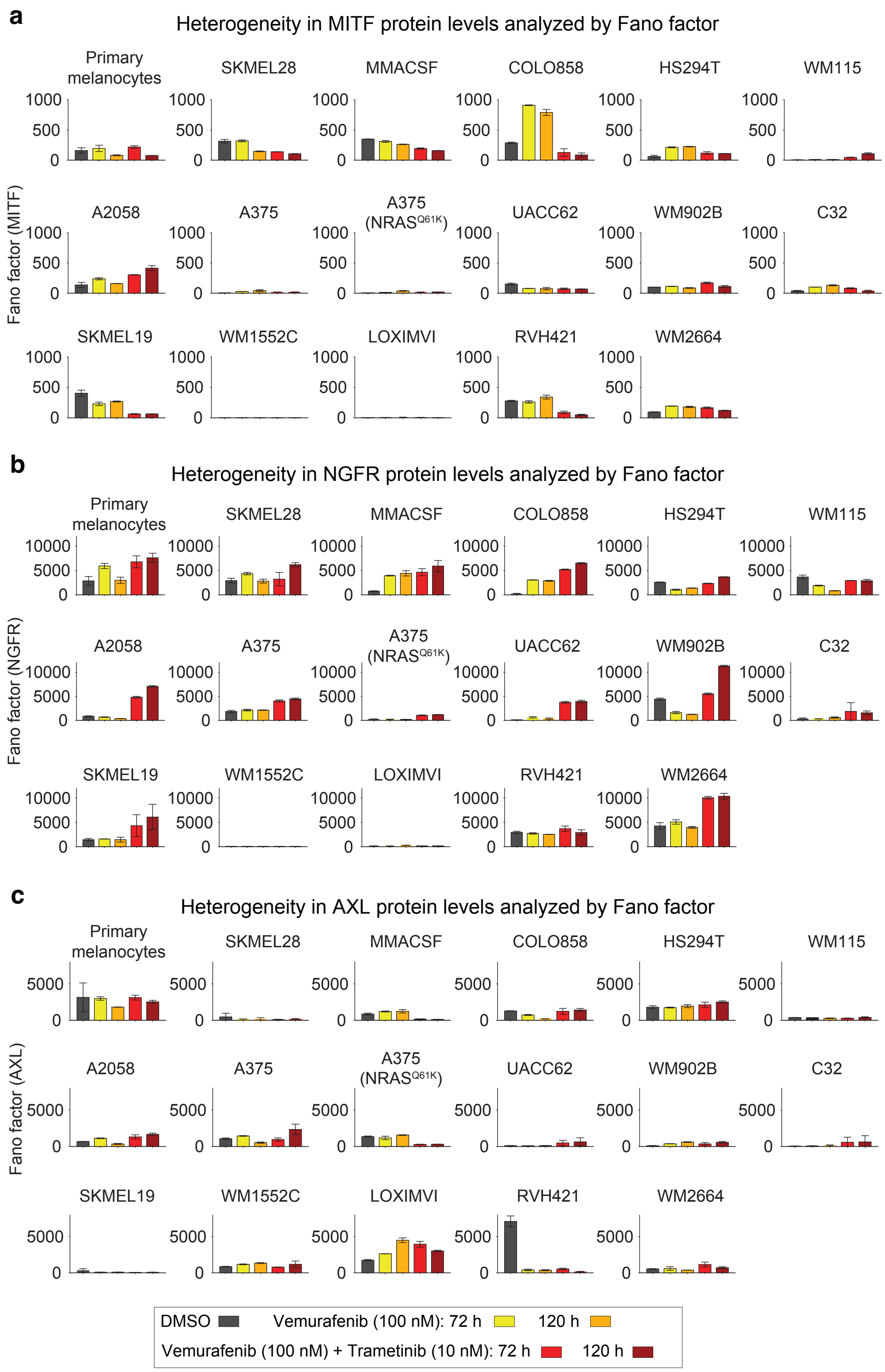


**Supplementary Figure 3. Heterogeneity in differentiation state markers MITF, NGFR and AXL, revealed by Fano factor analysis of primary melanocytes and 16 melanoma cell lines treated with BRAF/MEK inhibitors at indicated doses and timepoints.** Data represent mean values ± s.d. calculated across 2 replicates.

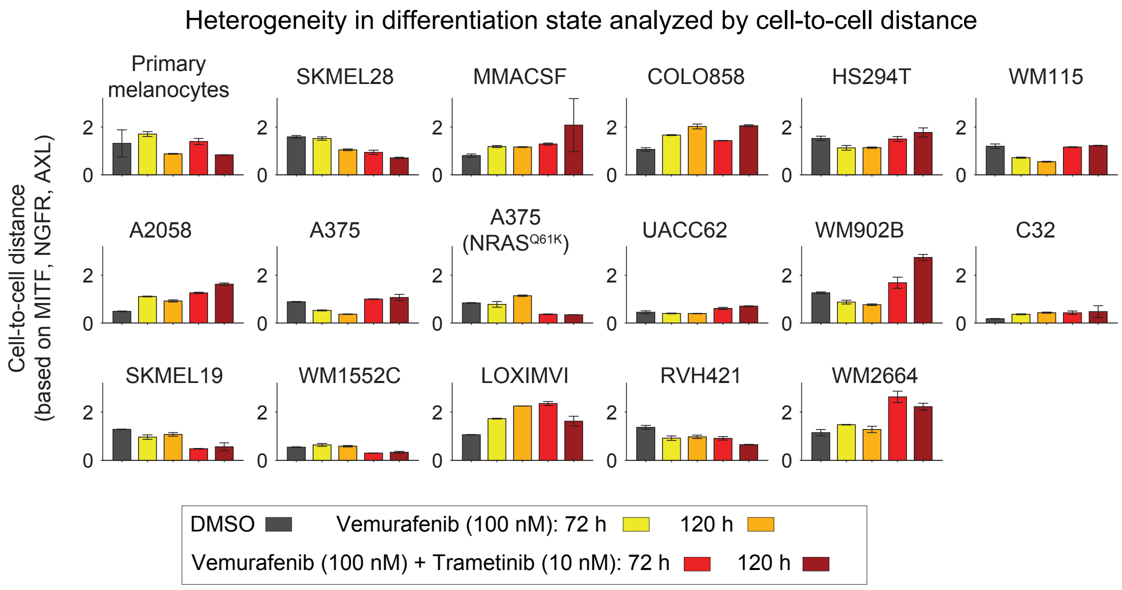


**Supplementary Figure 4. Heterogeneity in differentiation state markers MITF, NGFR and AXL, revealed by cell-to-cell distance analysis of primary melanocytes and 16 melanoma cell lines co-stained simultaneously for all three protein markers following treatment with BRAF/MEK inhibitors at indicated doses and timepoints.** Data represent mean values ± s.d. calculated across 2 replicates.

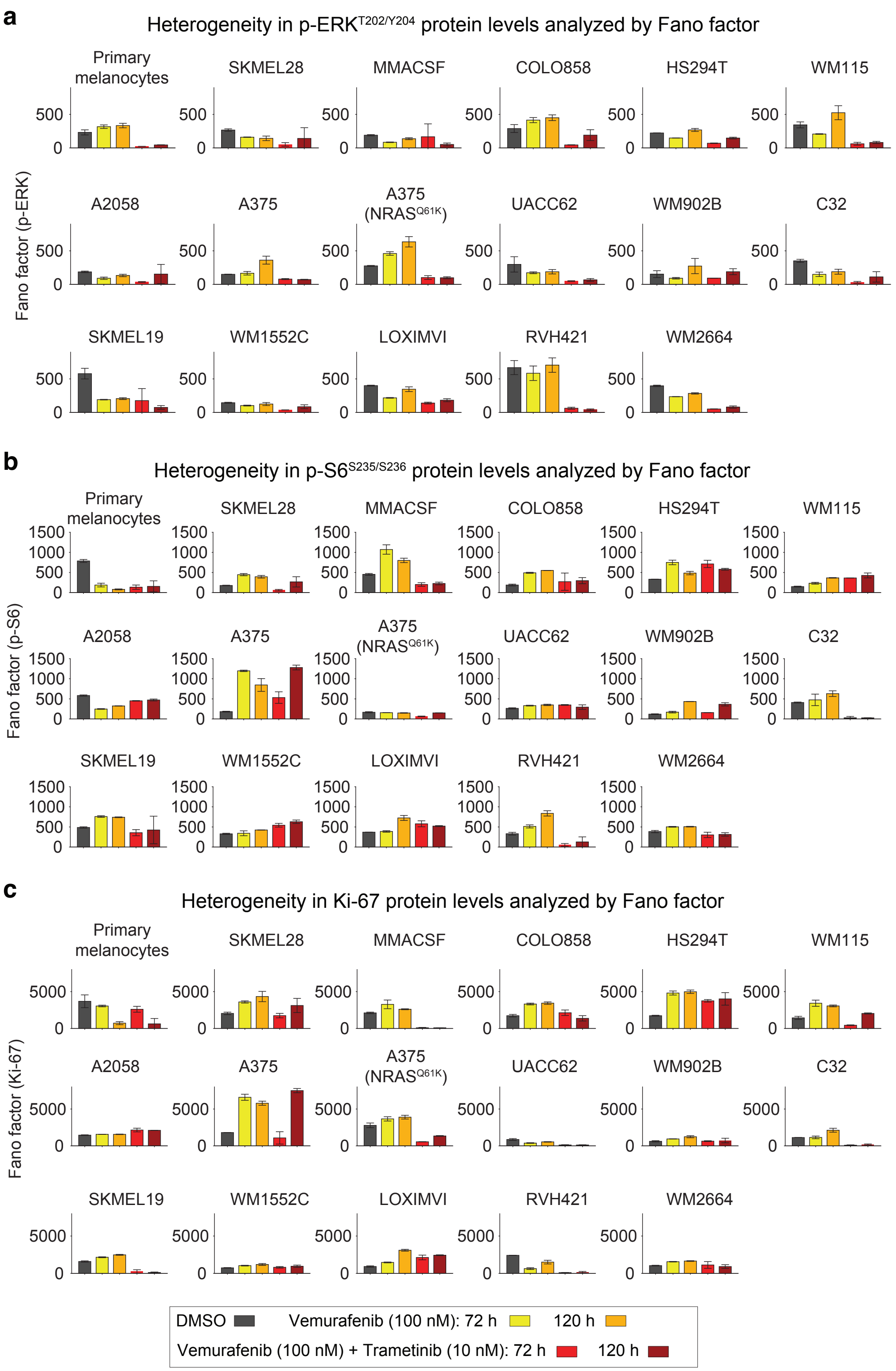


**Supplementary Figure 5. Heterogeneity in p-ERK$^{T202/Y204}$, p-S6$^{S235/S236}$ and Ki-67 proteins, revealed by Fano factor analysis of primary melanocytes and 16 melanoma cell lines treated with BRAF/MEK inhibitors at indicated doses and timepoints.** Data represent mean values ± s.d. calculated across 2 replicates.

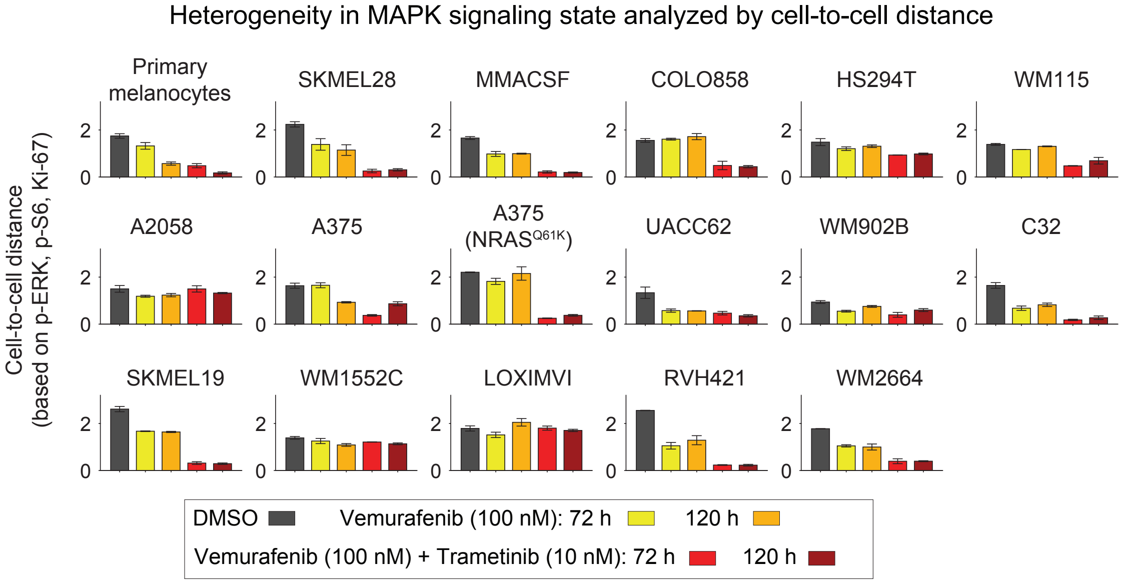


**Supplementary Figure 6. Heterogeneity in p-ERK$^{T202/Y204}$, p-S6$^{S235/S236}$ and Ki-67 proteins, revealed by cell-to-cell distance analysis of primary melanocytes and 16 melanoma cell lines co-stained simultaneously for all three protein markers following treatment with BRAF/MEK inhibitors at indicated doses and timepoints.** Data represent mean values ± s.d. calculated across 2 replicates.

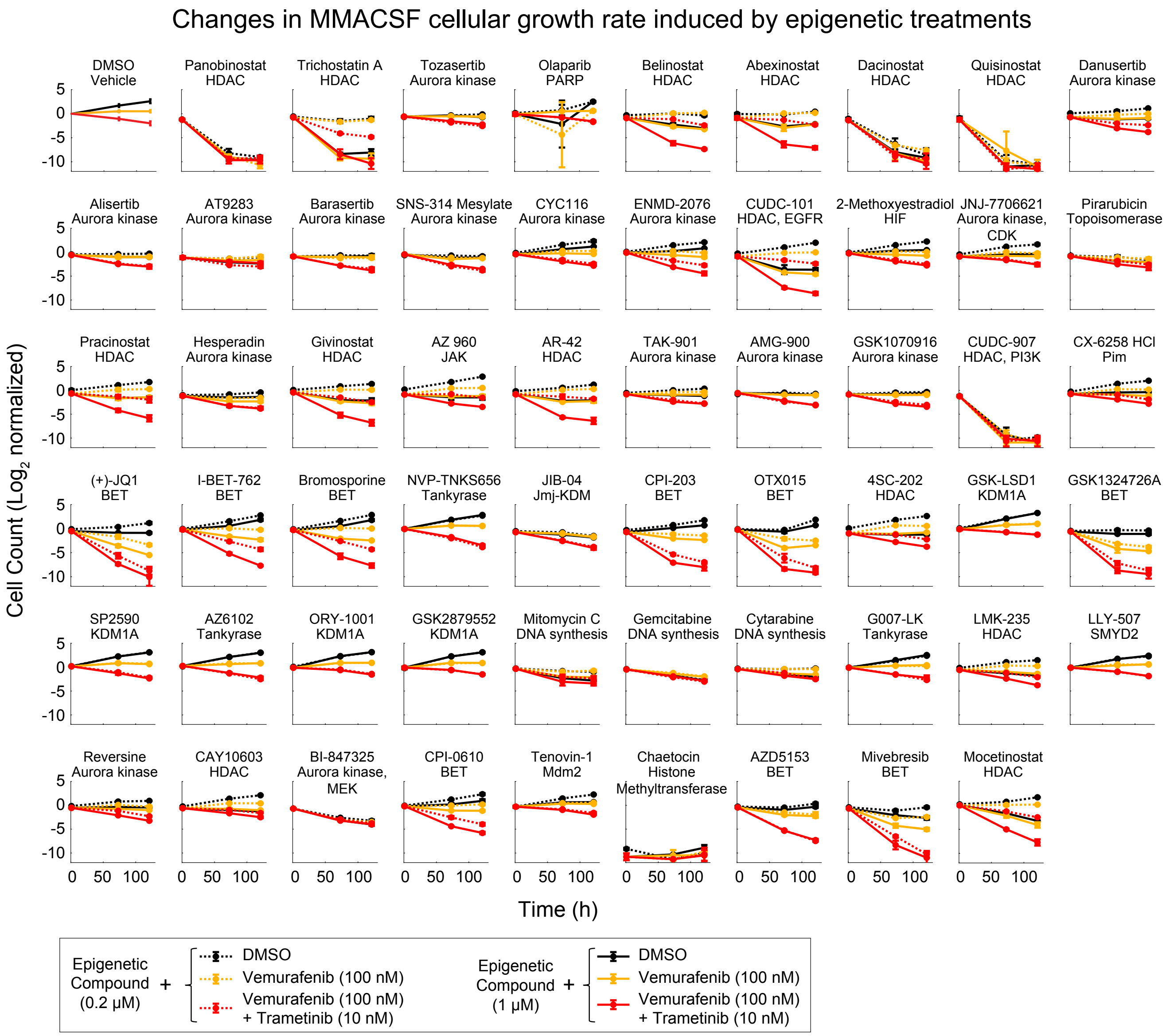


**Supplementary Figure 7. Changes in MMACSF cellular growth rate induced by significant epigenetic treatments.** $Log_2$-normalized changes in live cell count following exposure of MMACSF cells (pretreated for 24 h with either DMSO or two different doses of 58 significant epigenetic compounds) to either DMSO, vemurafenib (at 100 nM), or vemurafenib (at 100 nM ) plus trametinib (at 10 nM), for a period of 3-5 days. Individual compounds and their nominal epigenetic targets are indicated. Data represent mean values ± s.d. calculated for two replicates per treatment condition.

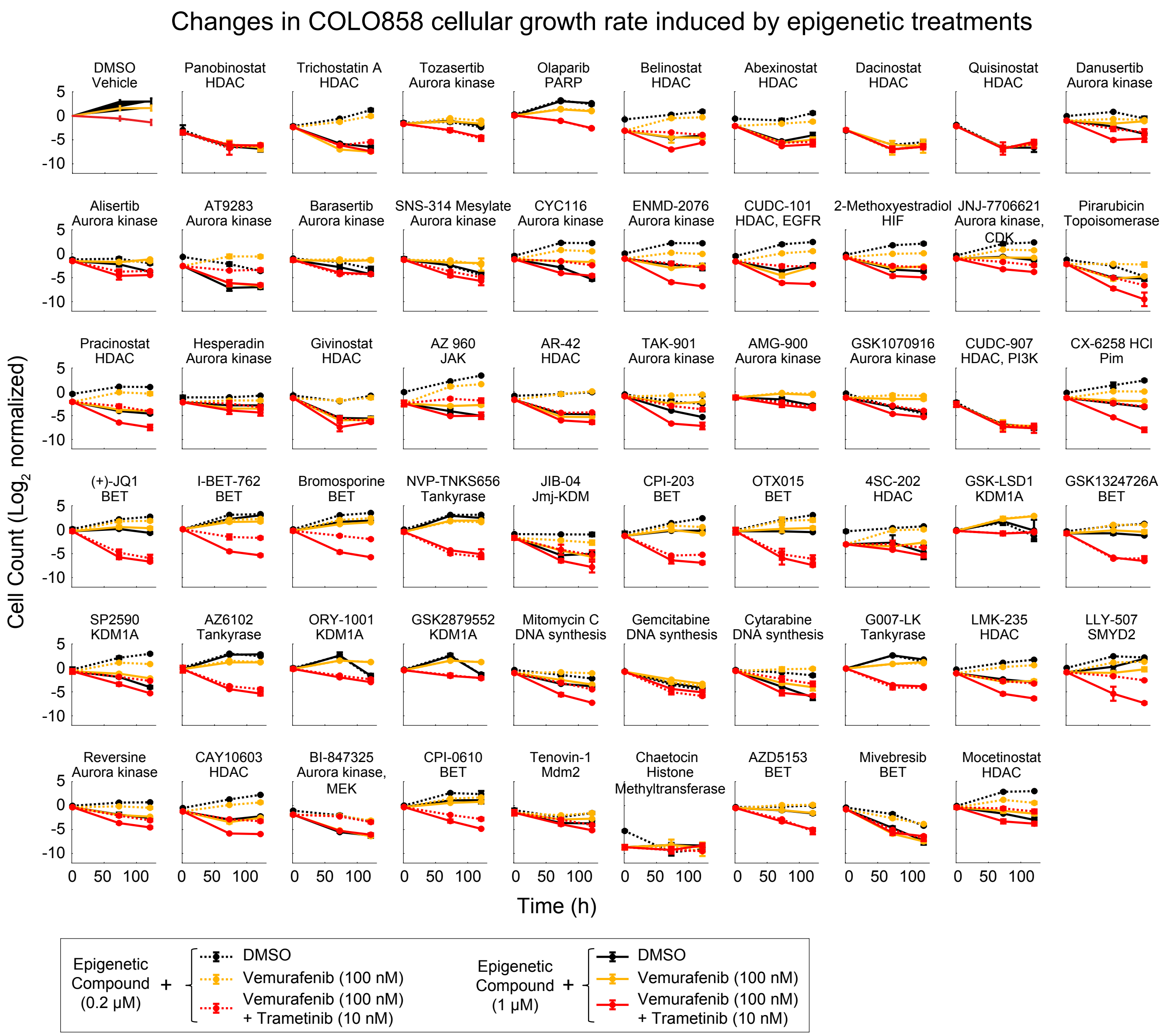


**Supplementary Figure 8. Changes in COLO858 cellular growth rate induced by significant epigenetic treatments.** $Log_2$-normalized changes in live cell count following exposure of COLO858 cells (pretreated for 24 h with either DMSO or two different doses of 58 significant epigenetic compounds) to either DMSO, vemurafenib (at 100 nM), or vemurafenib (at 100 nM ) plus trametinib (at 10 nM), for a period of 3-5 days. Individual compounds and their nominal epigenetic targets are indicated. Data represent mean values ± s.d. calculated for two replicates per treatment condition.

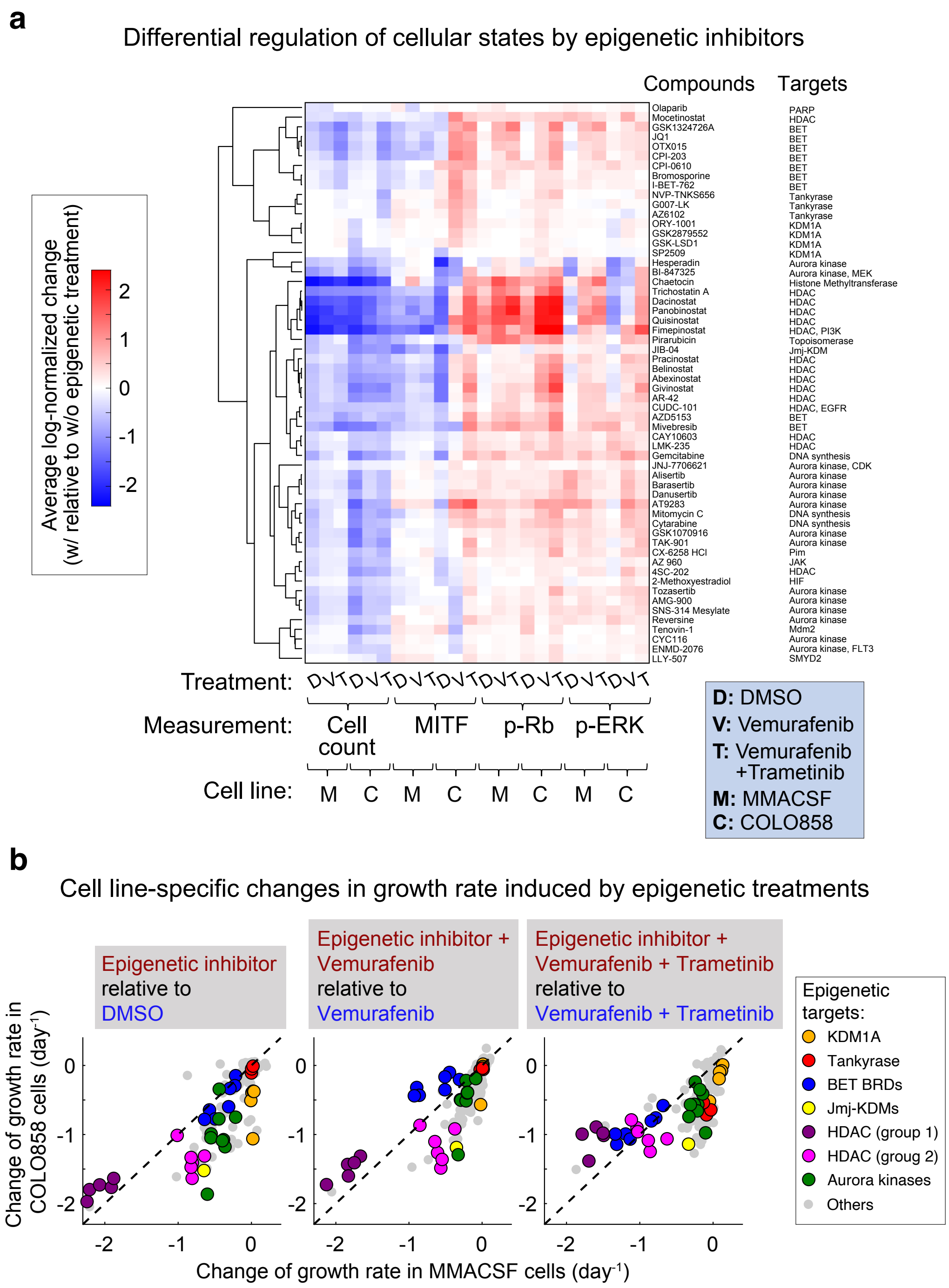


**Supplementary Figure 9. Differential regulation of cellular states by mechanistically distinct epigenetic-modifying compounds. (a)** Hierarchical clustering of normalized measurements of cellular grow rate, MITF, p-Rb$^{S807/811}$, and p-ERK$^{T202/Y204}$ at indicated treatment conditions and timepoints in MMACSF and COLO858 cells as described in Figure 2b. **(b)** Cell line-specific changes in growth rate induced by each class of epigenetic treatments. Growth rate data collected for cells treated with each epigenetic compound and MAPK inhibitor condition (i.e., DMSO, vemurafenib, and vemurafenib plus trametinib) were normalized to cells treated without any epigenetic compound and the same MAPK inhibitor condition.

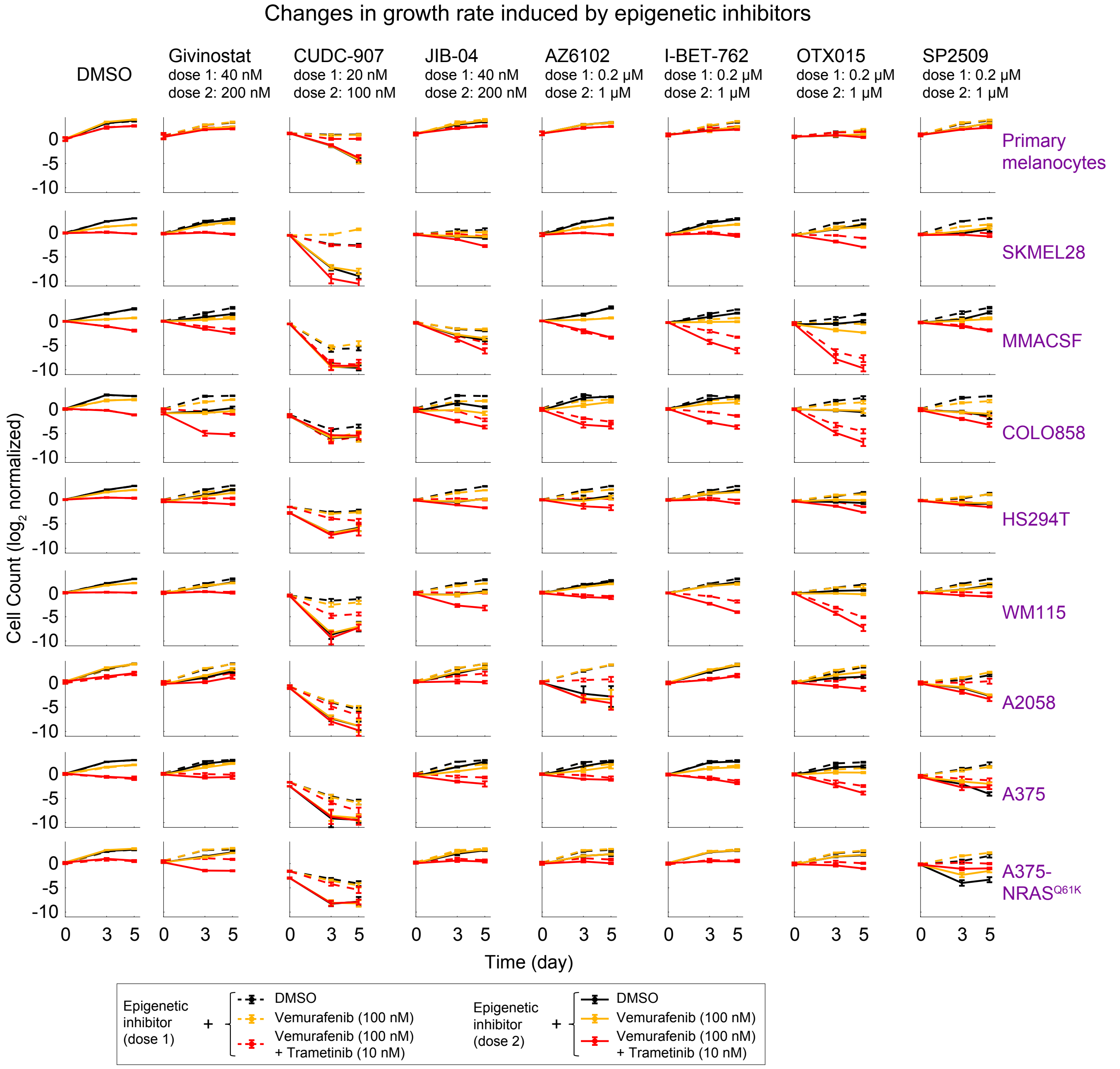


**Supplementary Figure 10. Epigenetic treatment-induced changes in growth rates of 8 melanoma cell lines and primary melanocytes to identify optimized drug doses.** $Log_2$-normalized changes in live cell count following exposure of eight different melanoma cell lines and non-transformed primary melanocytes (pretreated for 24 h with either DMSO or two different doses of seven different epigenetic compounds) to either DMSO, vemurafenib (at 100 nM), or vemurafenib (at 100 nM ) plus trametinib (at 10 nM), for a period of 3-5 days. Treatment doses for each compound are as follows: Givinostat (40 and 200 nM), CUDC-907 (20 and 100 nM), JIB-04 (40 and 200 nM), AZ6102 (0.2 and 1 μM), I-BET-762 (0.2 and 1 μM), OTX015 (0.2 and 1 μM), SP2509 (0.2 and 1 μM). Data represent mean values ± s.d. calculated for two replicates per treatment condition.

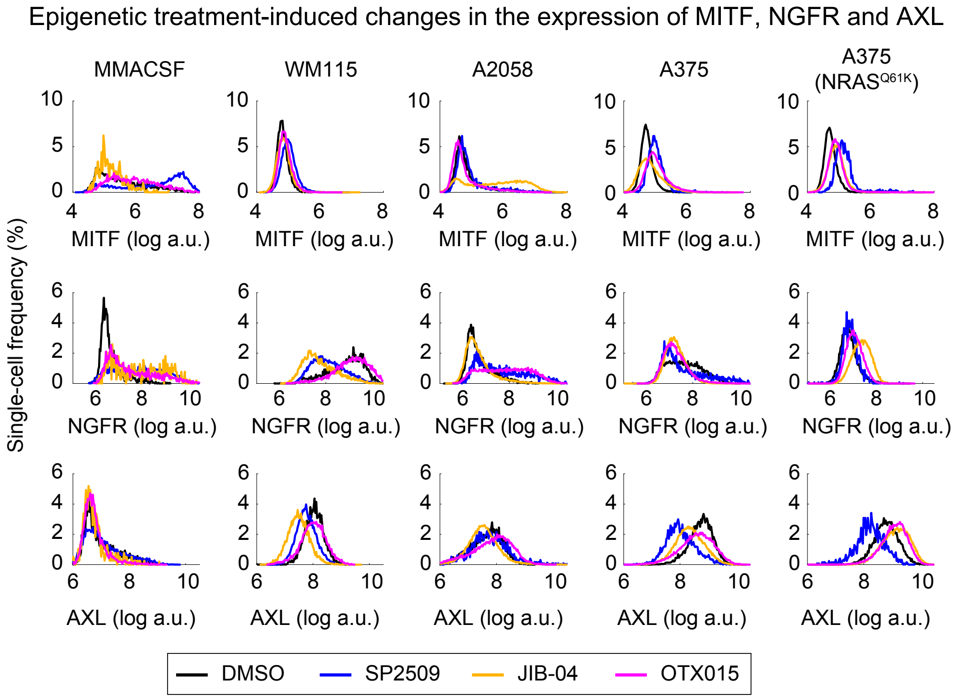


**Supplementary Figure 11. Single-cell protein levels of MITF (top), NGFR (middle) and AXL (bottom) measured by immunofluorescence microscopy, in five melanoma cell lines following treatment with vehicle (DMSO), SP2509, JIB-04 or OTX015 for 120 h.**

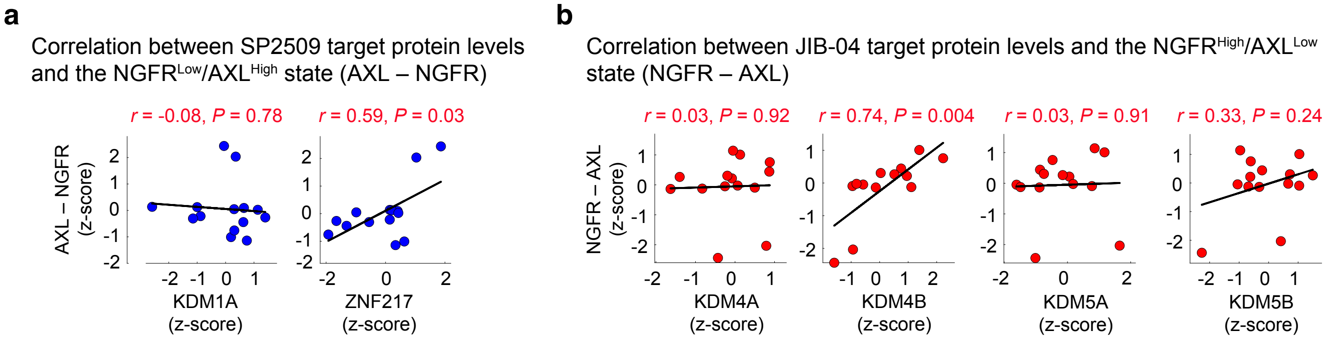


**Supplementary Figure 12. KDM4B and ZNF217 protein levels are correlated with relative levels of melanoma differentiation state markers NGFR and AXL. (a)** Pairwise Pearson correlation between variations in the difference between NGFR and AXL protein levels (NGFR – AXL or AXL – NGFR) and KDM1A (left) or ZNF217 (right) across BRAF-mutant melanoma cell lines. Protein data are extracted from the Cancer Cell Line Encyclopedia (CCLE) proteomics database (measured by multiplexed mass spectrometry) and z-scored across all of BRAF-mutant melanoma cell lines present in the database. **(b)** Pairwise Pearson correlation between variations in the difference between NGFR and AXL protein levels and JIB-04 targets, including KDM4A, KDM4B, KDM5A and KDM5B across BRAF-mutant melanoma cell lines. Protein data are extracted from the Cancer Cell Line Encyclopedia (CCLE) proteomics database (measured by multiplexed mass spectrometry) and z-scored across all of BRAF-mutant melanoma cell lines present in the database.

**Supplementary Table 1. List of chemicals used in this study, including their sources and catalog numbers, nominal targets, and purity.**

| Vendor | Cat # | Compound | Nominal target | Purity |
|---|---|---|---|---|
| Selleck Chemicals | S1004 | Veliparib (ABT-888) | PARP | 99.72% |
| Selleck Chemicals | S1007 | Roxadustat (FG-4592) | HIF | 99.39% |
| Selleck Chemicals | S1030 | Panobinostat (LBH589) | HDAC | 99.51% |
| Selleck Chemicals | S1045 | Trichostatin A (TSA) | HDAC | 97.62% |
| Selleck Chemicals | S1047 | Vorinostat (SAHA, MK0683) | Autophagy,HDAC | 99.90% |
| Selleck Chemicals | S1048 | Tozasertib (VX-680, MK-0457) | Aurora Kinase | 99.88% |
| Selleck Chemicals | S1053 | Entinostat (MS-275) | HDAC | 99.96% |
| Selleck Chemicals | S1060 | Olaparib (AZD2281, Ku-0059436) | PARP | 99.70% |
| Selleck Chemicals | S1085 | Belinostat (PXD101) | HDAC | 99.74% |
| Selleck Chemicals | S1087 | Iniparib (BSI-201) | PARP | 99.80% |
| Selleck Chemicals | S1090 | Abexinostat (PCI-24781) | HDAC | 97.29% |
| Selleck Chemicals | S1095 | Dacinostat (LAQ824) | HDAC | 95.40% |
| Selleck Chemicals | S1096 | Quisinostat (JNJ-26481585) 2HCl | HDAC | 99.62% |
| Selleck Chemicals | S1098 | Rucaparib (AG-014699,PF-01367338) phosphate | PARP | 99.89% |
| Selleck Chemicals | S1100 | MLN8054 | Aurora Kinase | 97.90% |
| Selleck Chemicals | S1103 | ZM 447439 | Aurora Kinase | 96.53% |
| Selleck Chemicals | S1107 | Danusertib (PHA-739358) | Aurora Kinase,Bcr-Abl,c-RET,FGFR | 99.55% |
| Selleck Chemicals | S1122 | Mocetinostat (MGCD0103) | HDAC | 99.46% |
| Selleck Chemicals | S1129 | SRT1720 HCl | Sirtuin | 99.07% |
| Selleck Chemicals | S1132 | INO-1001 (3-Aminobenzamide) | PARP | 99.80% |
| Selleck Chemicals | S1133 | Alisertib (MLN8237) | Aurora Kinase | 99.64% |
| Selleck Chemicals | S1134 | AT9283 | Aurora Kinase,Bcr-Abl,JAK | 100% |
| Selleck Chemicals | S1143 | AG-490 (Tyrphostin B42) | EGFR,JAK | 99.76% |
| Selleck Chemicals | S1147 | Barasertib (AZD1152-HQPA) | Aurora Kinase | 99.98% |
| Selleck Chemicals | S1154 | SNS-314 Mesylate | Aurora Kinase | 99.88% |
| Selleck Chemicals | S1168 | Valproic acid sodium salt (Sodium valproate) | GABA Receptor,HDAC,Autophagy | 100% |
| Selleck Chemicals | S1171 | CYC116 | Aurora Kinase,VEGFR | 99.52% |
| Selleck Chemicals | S1181 | ENMD-2076 | Aurora Kinase,FLT3,VEGFR | 99.41% |
| Selleck Chemicals | S1194 | CUDC-101 | EGFR,HDAC,HER2 | 99.36% |
| Selleck Chemicals | S1200 | Decitabine | DNA Methyltransferase | 99.93% |
| Selleck Chemicals | S1216 | PFI-1 (PF-6405761) | Epigenetic Reader Domain | 99.07% |
| Selleck Chemicals | S1233 | 2-Methoxyestradiol (2-MeOE2) | HIF | 99.68% |
| Selleck Chemicals | S1249 | JNJ-7706621 | Aurora Kinase,CDK | 99.45% |
| Selleck Chemicals | S1327 | Ellagic acid | Topoisomerase | 99.64% |
| Selleck Chemicals | S1378 | Ruxolitinib (INCB018424) | JAK | 99.92% |
| Selleck Chemicals | S1393 | Pirarubicin | Topoisomerase | 99.82% |
| Selleck Chemicals | S1396 | Resveratrol | Autophagy,Sirtuin | 99.89% |
| Selleck Chemicals | S1422 | Droxinostat | HDAC | 96.15% |
| Selleck Chemicals | S1451 | Aurora A Inhibitor I | Aurora Kinase | 99.40% |
| Selleck Chemicals | S1454 | PHA-680632 | Aurora Kinase | 96.62% |
| Selleck Chemicals | S1463 | Ofloxacin | Topoisomerase | 99.91% |
| Selleck Chemicals | S1484 | MC1568 | HDAC | 96.74% |
| Selleck Chemicals | S1509 | Norfloxacin | Topoisomerase | 100% |
| Selleck Chemicals | S1515 | Pracinostat (SB939) | HDAC | 99.77% |
| Selleck Chemicals | S1529 | Hesperadin | Aurora Kinase | 99.09% |
| Selleck Chemicals | S1541 | Selisistat (EX 527) | Sirtuin | 99.78% |
| Selleck Chemicals | S1782 | Azacitidine | DNA Methyltransferase | 99.92% |
| Selleck Chemicals | S2012 | PCI-34051 | HDAC | 99.01% |
| Selleck Chemicals | S2018 | ENMD-2076 L-(+)-Tartaric acid | Aurora Kinase,FLT3,VEGFR | 99.36% |
| Selleck Chemicals | S2158 | KW-2449 | Aurora Kinase,Bcr-Abl,FLT3 | 99.72% |
| Selleck Chemicals | S2162 | AZD1480 | JAK | 99.64% |
| Selleck Chemicals | S2170 | Givinostat (ITF2357) | HDAC | 99.07% |
| Selleck Chemicals | S2178 | AG-14361 | PARP | 99.65% |
| Selleck Chemicals | S2179 | Gandotinib (LY2784544) | JAK | 99.60% |
| Selleck Chemicals | S2198 | SGI-1776 free base | Pim | 99.39% |
| Selleck Chemicals | S2214 | AZ 960 | JAK | 99.05% |
| Selleck Chemicals | S2219 | Momelotinib (CYT387) | JAK | 99.08% |
| Selleck Chemicals | S2244 | AR-42 | HDAC | 99.03% |
| Selleck Chemicals | S2391 | Quercetin | Src,Sirtuin,PKC,PI3K | 99.60% |
| Selleck Chemicals | S2554 | Daphnetin | PKA,EGFR,PKC | 99.23% |
| Selleck Chemicals | S2627 | Tubastatin A HCl | HDAC | 99.30% |
| Selleck Chemicals | S2686 | NVP-BSK805 2HCl | JAK | 98.32% |
| Selleck Chemicals | S2692 | TG101209 | c-RET,FLT3,JAK | 99.74% |
| Selleck Chemicals | S2693 | Resminostat | HDAC | 99.04% |
| Selleck Chemicals | S2718 | TAK-901 | Aurora Kinase | 99.05% |
| Selleck Chemicals | S2719 | AMG-900 | Aurora Kinase | 99.03% |
| Selleck Chemicals | S2736 | Fedratinib (SAR302503, TG101348) | JAK | 99.96% |

| | | | | |
|---|---|---|---|---|
| Selleck Chemicals | S2740 | GSK1070916 | Aurora Kinase | 98.62% |
| Selleck Chemicals | S2759 | CUDC-907 | HDAC,PI3K | 99.38% |
| Selleck Chemicals | S2770 | MK-5108 (VX-689) | Aurora Kinase | 98.40% |
| Selleck Chemicals | S2779 | M344 | HDAC | 99.36% |
| Selleck Chemicals | S2789 | Tofacitinib (CP-690550,Tasocitinib) | JAK | 99.79% |
| Selleck Chemicals | S2796 | WP1066 | JAK | 99.49% |
| Selleck Chemicals | S2804 | Sirtinol | Sirtuin | 97% |
| Selleck Chemicals | S2806 | CEP-33779 | JAK | 99.56% |
| Selleck Chemicals | S2818 | Tacedinaline (CI994) | HDAC | 99.74% |
| Selleck Chemicals | S2821 | RG108 | DNA Methyltransferase,Transferase | 99.74% |
| Selleck Chemicals | S2851 | Baricitinib (LY3009104, INCB028050) | JAK | 99.88% |
| Selleck Chemicals | S2867 | WHI-P154 | EGFR,JAK | 99.02% |
| Selleck Chemicals | S2886 | PJ34 | PARP | 99.25% |
| Selleck Chemicals | S2902 | S-Ruxolitinib (INCB018424) | JAK | 99.81% |
| Selleck Chemicals | S2919 | IOX2 | HIF | 99.57% |
| Selleck Chemicals | S3001 | Clevudine | DNA/RNA Synthesis | 99.96% |
| Selleck Chemicals | S3147 | Entacapone | Histone Methyltransferase | 99.57% |
| Selleck Chemicals | S4125 | Sodium Phenylbutyrate | HDAC | 99.86% |
| Selleck Chemicals | S4246 | Tranylcypromine (2-PCPA) HCl | MAO | 99.84% |
| Selleck Chemicals | S4294 | Procainamide HCl | DNA Methyltransferase,Sodium Channel | 99.09% |
| Selleck Chemicals | S5001 | Tofacitinib (CP-690550) Citrate | JAK | 99.89% |
| Selleck Chemicals | S7029 | AZD2461 | PARP | 99.29% |
| Selleck Chemicals | S7036 | XL019 | JAK | 99.01% |
| Selleck Chemicals | S7041 | CX-6258 HCl | Pim | 99.41% |
| Selleck Chemicals | S7062 | Pinometostat (EPZ5676) | Histone Methyltransferase | 99.75% |
| Selleck Chemicals | S7070 | GSK J4 HCl | Histone Demethylase | 99.40% |
| Selleck Chemicals | S7079 | SGC 0946 | Histone Methyltransferase | 99.49% |
| Selleck Chemicals | S7088 | UNC1215 | Epigenetic Reader Domain | 99.73% |
| Selleck Chemicals | S7104 | AZD1208 | Pim | 99.86% |
| Selleck Chemicals | S7110 | (+)-JQ1 | Epigenetic Reader Domain | 99.84% |
| Selleck Chemicals | S7113 | Zebularine | DNA Methyltransferase | 99.67% |
| Selleck Chemicals | S7120 | 3-deazaneplanocin A (DZNeP) HCl | Histone Methyltransferase | 99.81% |
| Selleck Chemicals | S7152 | C646 | Histone Acetyltransferase | 98.73% |
| Selleck Chemicals | S7189 | I-BET-762 | Epigenetic Reader Domain | 99.05% |
| Selleck Chemicals | S7229 | RGFP966 | HDAC | 99.74% |
| Selleck Chemicals | S7231 | GSK2801 | Epigenetic Reader Domain | 99.06% |
| Selleck Chemicals | S7233 | Bromosporine | Epigenetic Reader Domain | 99.92% |
| Selleck Chemicals | S7234 | IOX1 | Histone Demethylase | 99.04% |
| Selleck Chemicals | S7237 | OG-L002 | Histone Demethylase | 99.38% |
| Selleck Chemicals | S7238 | NVP-TNKS656 | Tankyrase | 99.85% |
| Selleck Chemicals | S7256 | SGC-CBP30 | Epigenetic Reader Domain | 99.11% |
| Selleck Chemicals | S7265 | MM-102 | Histone Methyltransferase | 99.00% |
| Selleck Chemicals | S7276 | SGI-1027 | DNA Methyltransferase | 99.58% |
| Selleck Chemicals | S7281 | JIB-04 | Jumonji histone demethylases | 99.80% |
| Selleck Chemicals | S7292 | RG2833 (RGFP109) | HDAC | 99.17% |
| Selleck Chemicals | S7294 | PFI-2 HCl | Histone Methyltransferase | 99.05% |
| Selleck Chemicals | S7295 | RVX-208 | Epigenetic Reader Domain | 99.47% |
| Selleck Chemicals | S7296 | ML324 | Histone Demethylase | 99.02% |
| Selleck Chemicals | S7300 | PJ34 HCl | PARP | 99.25% |
| Selleck Chemicals | S7304 | CPI-203 | Epigenetic Reader Domain | 99.69% |
| Selleck Chemicals | S7305 | MS436 | Epigenetic Reader Domain | 95.98% |
| Selleck Chemicals | S7315 | PFI-3 | Epigenetic Reader Domain | 99.87% |
| Selleck Chemicals | S7324 | TMP269 | HDAC | 99.00% |
| Selleck Chemicals | S7353 | EPZ004777 | Histone Methyltransferase | 99.09% |
| Selleck Chemicals | S7360 | OTX015 | Epigenetic Reader Domain | 99.81% |
| Selleck Chemicals | S7373 | UNC669 | Epigenetic Reader Domain | 99.07% |
| Selleck Chemicals | S7438 | ME0328 | PARP | 99.89% |
| Selleck Chemicals | S7473 | Nexturastat A | HDAC | 99.65% |
| Selleck Chemicals | S7476 | MG149 | Histone Acetyltransferase | 99.52% |
| Selleck Chemicals | S7541 | Decernotinib (VX-509) | JAK | 99.57% |
| Selleck Chemicals | S7555 | 4SC-202 | HDAC | 99.84% |
| Selleck Chemicals | S7570 | UNC0379 | Histone Methyltransferase | 99.66% |
| Selleck Chemicals | S7572 | A-366 | Histone Methyltransferase | 99.23% |
| Selleck Chemicals | S7574 | GSK-LSD1 2HCl | Histone Demethylase KDM1A | 99.78% |
| Selleck Chemicals | S7581 | GSK J1 | Histone Demethylase | 99.67% |
| Selleck Chemicals | S7582 | Anacardic Acid | Histone Acetyltransferase | 99.72% |
| Selleck Chemicals | S7591 | BRD4770 | Histone Methyltransferase | 99.75% |
| Selleck Chemicals | S7605 | Filgotinib (GLPG0634) | JAK | 99.72% |
| Selleck Chemicals | S7610 | UNC0631 | Histone Methyltransferase | 99.35% |
| Selleck Chemicals | S7611 | EI1 | Histone Methyltransferase | 99.04% |

| | | | | |
|---|---|---|---|---|
| Selleck Chemicals | S7616 | CPI-169 | Histone Methyltransferase | 99.00% |
| Selleck Chemicals | S7618 | MI-2 (Menin-MLL Inhibitor) | Histone Methyltransferase | 99.39% |
| Selleck Chemicals | S7619 | MI-3 (Menin-MLL Inhibitor) | Histone Methyltransferase | 99.03% |
| Selleck Chemicals | S7620 | GSK1324726A (I-BET726) | Epigenetic Reader Domain | 99.61% |
| Selleck Chemicals | S7625 | Niraparib (MK-4827) tosylate | PARP | 99.69% |
| Selleck Chemicals | S7641 | Remodelin | Histone Acetyltransferase | 99.39% |
| Selleck Chemicals | S7656 | CPI-360 | Histone Methyltransferase | 99.72% |
| Selleck Chemicals | S7680 | SP2509 | Histone Demethylase KDM1A | 99.05% |
| Selleck Chemicals | S7681 | OF-1 | Epigenetic Reader Domain | 99.19% |
| Selleck Chemicals | S7748 | EPZ015666(GSK3235025) | Histone Methyltransferase | >97% |
| Selleck Chemicals | S7767 | AZ6102 | Tankyrase | 99.58% |
| Selleck Chemicals | S7795 | ORY-1001 (RG-6016) 2HCl | Histone Demethylase KDM1A | 99.95% |
| Selleck Chemicals | S7796 | GSK2879552 2HCl | Histone Demethylase KDM1A | 99.51% |
| Selleck Chemicals | S7804 | GSK503 | Histone Methyltransferase | 99.84% |
| Selleck Chemicals | S7805 | EPZ011989 | Histone Methyltransferase | 99.07% |
| Selleck Chemicals | S7832 | SGC707 | Histone Methyltransferase | 99.77% |
| Selleck Chemicals | S7835 | I-BRD9 | Epigenetic Reader Domain | 99.00% |
| Selleck Chemicals | S8001 | Ricolinostat (ACY-1215) | HDAC | 99.89% |
| Selleck Chemicals | S8004 | ZM 39923 HCl | JAK | 99.80% |
| Selleck Chemicals | S8005 | SMI-4a | Pim | 100.00% |
| Selleck Chemicals | S8006 | BIX 01294 | Histone Methyltransferase | 100.00% |
| Selleck Chemicals | S8038 | UPF 1069 | PARP | 99.47% |
| Selleck Chemicals | S8043 | Scriptaid | HDAC | 99.22% |
| Selleck Chemicals | S8049 | Tubastatin A | HDAC | 99.67% |
| Selleck Chemicals | S8056 | Lomeguatrib | DNA Methyltransferase | 99.37% |
| Selleck Chemicals | S8057 | Pacritinib (SB1518) | FLT3,JAK | 97.09% |
| Selleck Chemicals | S8096 | Mirin | ATM/ATR | 99.13% |
| Selleck Chemicals | S8111 | GSK591 | Histone Methyltransferase | 99.00% |
| Selleck Chemicals | S8112 | MS023 | Histone Methyltransferase | 99.15% |
| Selleck Chemicals | S8146 | Mitomycin C | DNA/RNA Synthesis | 99.74% |
| Selleck Chemicals | S8179 | BI-7273 | Epigenetic Reader Domain | 99.79% |
| Selleck Chemicals | S8180 | PF-CBP1 HCl | Epigenetic Reader Domain | 99.01% |
| Selleck Chemicals | S8195 | Oclacitinib | JAK | 99.09% |
| Selleck Chemicals | S8209 | HLCL-61 HCL | Histone Methyltransferase | 99.79% |
| Selleck Chemicals | S8323 | ITSA-1 (ITSA1) | HDAC | 99.08% |
| Selleck Chemicals | S8481 | SRT3025 HCl | Sirtuin | 99.62% |
| Selleck Chemicals | S8496 | EED226 | Epigenetic Reader Domain | 99.05% |
| Selleck Chemicals | S8502 | TMP195 | HDAC | 99.05% |
| Selleck Chemicals | S8567 | Tucidinostat (Chidamide) | HDAC | 99.09% |
| Selleck Chemicals | S1055 | Enzastaurin (LY317615) | PKC | 99.47% |
| Selleck Chemicals | S1573 | Fasudil (HA-1077) HCl | Autophagy,ROCK | 100% |
| Selleck Chemicals | S1703 | Divalproex Sodium | HDAC | 99.91% |
| Selleck Chemicals | S1774 | Thioguanine | DNA Methyltransferase | 99.93% |
| Selleck Chemicals | S1802 | AICAR (Acadesine) | AMPK | 99.97% |
| Selleck Chemicals | S1848 | Curcumin | NF-κB,HDAC,Histone Acetyltransferase,Nrf2 | 99.83% |
| Selleck Chemicals | S1899 | Nicotinamide (Vitamin B3) | Sirtuin | 100% |
| Selleck Chemicals | S2197 | A-966492 | PARP | 99.80% |
| Selleck Chemicals | S2250 | (-)-Epigallocatechin Gallate | DNA Methyltransferase,HER2,Telomerase,EGFR,Fatty Acid Synthase | 99.68% |
| Selleck Chemicals | S2298 | Fisetin | Sirtuin | 97.76% |
| Selleck Chemicals | S2341 | (-)-Parthenolide | HDAC,NF-κB,Mdm2,p53 | 100% |
| Selleck Chemicals | S2407 | Curcumol | JAK | >95% |
| Selleck Chemicals | S2542 | Phenformin HCl | AMPK | 99.05% |
| Selleck Chemicals | S2697 | A-769662 | AMPK,Fatty Acid Synthase | 99.12% |
| Selleck Chemicals | S2791 | Sotrastaurin | PKC | 99.57% |
| Selleck Chemicals | S2911 | Go 6983 | PKC | 97.25% |
| Selleck Chemicals | S4170 | Coumarin | Immunology & Inflammation related | 99.97% |
| Selleck Chemicals | S4589 | Amodiaquine dihydrochloride dihydrate | Transferase,Histone Methyltransferase | 99.80% |
| Selleck Chemicals | S4710 | Picolinamide | PARP | 99.93% |
| Selleck Chemicals | S4715 | Benzamide | PARP | 98.15% |
| Selleck Chemicals | S4735 | Salvianolic acid B | Sirtuin | 99.73% |
| Selleck Chemicals | S4900 | Tenovin-6 | p53,Sirtuin | 98.61% |
| Selleck Chemicals | S7065 | MK-8745 | Aurora Kinase | 99.05% |
| Selleck Chemicals | S7119 | Go6976 | FLT3,JAK,PKC | 99.34% |
| Selleck Chemicals | S7128 | Tazemetostat (EPZ-6438) | Histone Methyltransferase | 99.56% |
| Selleck Chemicals | S7144 | BMS-911543 | JAK | 99.03% |
| Selleck Chemicals | S7165 | UNC1999 | Histone Methyltransferase | 99.20% |
| Selleck Chemicals | S7207 | Bisindolylmaleimide IX (Ro 31-8220 Mesylate) | PKC | 99.02% |
| Selleck Chemicals | S7239 | G007-LK | Tankyrase | 99.00% |
| Selleck Chemicals | S7259 | FLLL32 | JAK | 95.54% |
| Selleck Chemicals | S7278 | HPOB | HDAC | 99.36% |

| | | | | |
|---|---|---|---|---|
| Selleck Chemicals | S7317 | WZ4003 | AMPK | 98.00% |
| Selleck Chemicals | S7318 | HTH-01-015 | AMPK | 99.07% |
| Selleck Chemicals | S7569 | LMK-235 | HDAC | 99.05% |
| Selleck Chemicals | S7575 | LLY-507 | Histone Methyltransferase | 97.33% |
| Selleck Chemicals | S7577 | AGK2 | Sirtuin | 99.07% |
| Selleck Chemicals | S7588 | Reversine | Adenosine Receptor,Aurora Kinase | 96.15% |
| Selleck Chemicals | S7593 | Splitomicin | HDAC | 96.04% |
| Selleck Chemicals | S7595 | Santacruzamate A (CAY10683) | HDAC | 99.57% |
| Selleck Chemicals | S7596 | CAY10603 | HDAC | 99.04% |
| Selleck Chemicals | S7612 | PX-478 2HCl | HIF | >97% |
| Selleck Chemicals | S7617 | Tasquinimod | HDAC | 99.29% |
| Selleck Chemicals | S7634 | Cerdulatinib (PRT062070, PRT2070) | JAK | 97.82% |
| Selleck Chemicals | S7650 | Peficitinib (ASP015K, JNJ-54781532) | JAK | 99.67% |
| Selleck Chemicals | S7689 | BG45 | HDAC | 97.91% |
| Selleck Chemicals | S7726 | BRD73954 | HDAC | 99.65% |
| Selleck Chemicals | S7730 | NU1025 | PARP | 99.29% |
| Selleck Chemicals | S7792 | SRT2104 (GSK2245840) | Sirtuin | 99.61% |
| Selleck Chemicals | S7815 | MI-136 | Histone Methyltransferase | 99.05% |
| Selleck Chemicals | S7816 | MI-463 | Histone Methyltransferase | 99.35% |
| Selleck Chemicals | S7817 | MI-503 | Histone Methyltransferase | 99.16% |
| Selleck Chemicals | S7820 | EPZ020411 2HCl | Histone Methyltransferase | 99.08% |
| Selleck Chemicals | S7833 | OICR-9429 | Histone Methyltransferase | 99.14% |
| Selleck Chemicals | S7843 | BI-847325 | MEK,Aurora Kinase | 95.94% |
| Selleck Chemicals | S7845 | SirReal2 | Sirtuin | 99.50% |
| Selleck Chemicals | S7853 | CPI-0610 | Epigenetic Reader Domain | 99.95% |
| Selleck Chemicals | S7884 | AMI-1 | Histone Methyltransferase | 99.02% |
| Selleck Chemicals | S7906 | PFI-4 | Epigenetic Reader Domain | 99.74% |
| Selleck Chemicals | S7946 | KC7F2 | HIF | 99.03% |
| Selleck Chemicals | S7953 | ETC-1002 | AMPK,LDL | 99.13% |
| Selleck Chemicals | S7958 | Lificiguat(YC-1) | HIF | 99.94% |
| Selleck Chemicals | S7979 | FG-2216 | HIF | 99.78% |
| Selleck Chemicals | S7983 | A-196 | Histone Methyltransferase | 99.53% |
| Selleck Chemicals | S8000 | Tenovin-1 | E3 Ligase ,p53 | 99.82% |
| Selleck Chemicals | S8068 | Chaetocin | Histone Methyltransferase | 98.34% |
| Selleck Chemicals | S8071 | UNC0638 | Histone Methyltransferase | 99.08% |
| Selleck Chemicals | S8138 | Molidustat (BAY 85-3934) | HIF | 99.82% |
| Selleck Chemicals | S8147 | MS049 | Histone Methyltransferase | 99.48% |
| Selleck Chemicals | S8171 | Daprodustat (GSK1278863) | HIF | 99.29% |
| Selleck Chemicals | S8190 | CPI-637 | Epigenetic Reader Domain | 99.61% |
| Selleck Chemicals | S8245 | Thiomyristoyl | Sirtuin | 99.84% |
| Selleck Chemicals | S8249 | HPI-4 (Ciliobrevin A) | Hedgehog/Smoothened | 99.63% |
| Selleck Chemicals | S8265 | GSK6853 | Epigenetic Reader Domain | 99.44% |
| Selleck Chemicals | S8270 | SRT2183 | Sirtuin | 99.19% |
| Selleck Chemicals | S8287 | CPI-455 HCl | Histone Demethylase | 99.08% |
| Selleck Chemicals | S8340 | SGC2085 | Histone Methyltransferase | 99.05% |
| Selleck Chemicals | S8344 | AZD5153 | Epigenetic Reader Domain | 98.19% |
| Selleck Chemicals | S8353 | CPI-1205 | Histone Methyltransferase | 99.04% |
| Selleck Chemicals | S8359 | UNC3866 | Histone Methyltransferase | 96.46% |
| Selleck Chemicals | S8363 | NMS-P118 | PARP | 99.90% |
| Selleck Chemicals | S8370 | BGP-15 2HCl | PARP | 99.94% |
| Selleck Chemicals | S8400 | Mivebresib(ABBV-075) | Epigenetic Reader Domain | 99.23% |
| Selleck Chemicals | S8419 | E7449 | PARP | 97.48% |
| Selleck Chemicals | S8429 | PNU-74654 | Wnt/beta-catenin | 99.49% |
| Selleck Chemicals | S8441 | LW 6 | HIF | 99.00% |
| Selleck Chemicals | S8443 | MK-8617 | HIF | 99.39% |
| Selleck Chemicals | S8460 | Salermide | Sirtuin | 99.79% |
| Selleck Chemicals | S8464 | Citarinostat (ACY-241) | HDAC | 99.03% |
| Selleck Chemicals | S1149 | Gemcitabine HCl | Autophagy,DNA/RNA Synthesis | 99.96% |
| Selleck Chemicals | S1215 | Carboplatin | DNA/RNA Synthesis | 99.26% |
| Selleck Chemicals | S1373 | Daptomycin | DNA/RNA Synthesis | 99.30% |
| Selleck Chemicals | S1384 | Mizoribine | DNA/RNA Synthesis | 99.96% |
| Selleck Chemicals | S1648 | Cytarabine | DNA/RNA Synthesis | 99.98% |
| Selleck Chemicals | S1826 | Nedaplatin | DNA/RNA Synthesis | 99.74% |
| Selleck Chemicals | S1995 | Procarbazine HCl | DNA/RNA Synthesis | 99.01% |
| Selleck Chemicals | S7419 | Blasticidin S HCl | DNA/RNA Synthesis | 99.91% |
| Selleck Chemicals | S8197 | APTSTAT3-9R | STAT | 99.56% |
| Selleck Chemicals | S1950 | Metformin HCl | Autophagy | 99.87% |
| Selleck Chemicals | S1999 | Sodium butyrate | HDAC | 100% |
| Selleck Chemicals | S7306 | Dorsomorphin (Compound C) 2HCl | AMPK | 99.67% |
| Selleck Chemicals | S1267 | Vemurafenib | BRAF | 99.26% |

| | | | | |
|---|---|---|---|---|
| Selleck Chemicals | S2673 | Trametinib | MEK | 99.65% |
| Medchem Express | HY-103713 | Seclidemstat (SP-2577) | Histone Demethylase KDM1A | 98.78% |

**Supplementary Table 2. Small molecule screening data**

| Category | Parameter | Description |
|---|---|---|
| Assay | Type of assay | In vitro phenotypic profiling of small molecule epigenetic-modifying compound's effects (individually or in combination with BRAF/MEK inhibitors) on melanoma cells |
| | Target | Epigenetic modifiers |
| | Primary measurement | Melanoma cell count (by DAPI) using fluorescence microscopy |
| | Key reagents | See chemical reagent table and methods |
| | Assay protocol | "Epigenetic compound screen" and "Immunofluorescence staining, quantitation and analysis" in manuscript |
| | Additional comments | |
| Library | Library size | 276 |
| | Library composition | Epigenetic-modifying compounds |
| | Source | Selleck Chemicals |
| | Additional comments | |
| Screen | Format | 96-well plates |
| | Concentration(s) tested | 0.2 µM and 1 µM for all compounds; equivalent volume of DMSO or water as vehicle control |
| | Plate controls | DMSO or water |
| | Reagent/ compound dispensing system | HP D300e Digital Dispenser |
| | Detection instrument and software | ImageXpress Micro Confocal High-Content Imaging System (Molecular Devices) and MetaXpress Imaging software, version 6.2.3.733 |
| | Assay validation/QC | Replicate $R^2 > 0.98$ |
| | Correction factors | |
| | Normalization | Growth rate, MITF, p-Rb and p-ERK levels for cells treated with each epigenetic compound were normalized to cells treated with the vehicle (DMSO or water). |
| | Additional comments | |
| Post-HTS analysis | Hit criteria | A statistically significant decrease in normalized growth rate of cells treated with the epigenetic compound in at least one of the two cell lines and at least one of the tested MAPK inhibitor conditions (vemurafenib, vemurafenib plus trametinib or DMSO vehicle) |
| | Hit rate | 58/276 ≈ 21% |
| | Additional assay(s) | Immunofluorescence microscopy of cell signaling (p-ERK), differentiation state (MITF) and proliferation marker (p-Rb) |
| | Confirmation of hit purity and structure | N/A |
| | Additional comments | Screen data made available |

**Supplementary Dataset 1.** Single-cell protein levels of three melanoma differentiation state markers, including MITF, NGFR, and AXL, measured by multiplexed immunofluorescence microscopy across 16 $BRAF^{V600E/D}$ melanoma cell lines and primary melanocytes following exposure to either vehicle (DMSO), BRAF inhibitor (vemurafenib at 100 nM), or the combination of BRAF and MEK inhibitors (vemurafenib at 100 nM and trametinib at 10 nM) for 3 and 5 days. Single-cell data for a maximum of 8,000 randomly selected cells per condition are shown. Data are associated with Fig. 1a, b.

**Supplementary Dataset 2.** Single-cell protein levels of MAPK signaling and proliferation state markers, including p-$ERK^{T202/Y204}$, p-$S6^{S235/S236}$, and Ki-67, measured by multiplexed immunofluorescence microscopy across 16 $BRAF^{V600E/D}$ melanoma cell lines and primary melanocytes following exposure to either vehicle (DMSO), BRAF inhibitor (vemurafenib at 100 nM), or the combination of BRAF and MEK inhibitors (vemurafenib at 100 nM and trametinib at 10 nM) for 3 and 5 days. Single-cell data for a maximum of 8,000 randomly selected cells per condition are shown. Data are associated with Fig. 1c, d.

**Supplementary Dataset 3.** Measurements of live cell count for 16 $BRAF^{V600E/D}$ melanoma cell lines and primary melanocytes following exposure to either vehicle (DMSO), BRAF inhibitor (vemurafenib at 100 nM), or the combination of BRAF and MEK inhibitors (vemurafenib at 100 nM and trametinib at 10 nM) for 3 and 5 days. Data are associated with Fig. 1e, f.

**Supplementary Dataset 4.** Measurements of live cell count following exposure of COLO858 and MMACSF cells to either DMSO, vemurafenib (at 100 nM), or vemurafenib (at 100 nM) plus trametinib (at 10 nM) for a period of 72 or 120 h following pretreatment for 24 h with either DMSO or two different doses (0.2 and 1 μM) of each of the 276 epigenetic-modifying compounds. Data are associated with Fig. 2b.

**Supplementary Dataset 5.** Measurements of live cell count, MITF, p-$Rb^{S807/811}$, and p-$ERK^{T202/Y204}$ following exposure of COLO858 and MMACSF cells to either DMSO, vemurafenib (at 100 nM), or vemurafenib (at 100 nM) plus trametinib (at 10 nM) for a period of 72 or 120 h following pretreatment for 24 h with either DMSO or two different doses (0.2 and 1 μM) of each of the 58 significant epigenetic-modifying compounds. Data are associated with Fig. 2c.

**Supplementary Dataset 6.** Measurements of live cell count following exposure of 16 melanoma cell lines and non-transformed primary melanocytes to either DMSO, vemurafenib (at 100 nM), or vemurafenib (at 100 nM) plus trametinib (at 10 nM) for a period of 72 or 120 h following pretreatment for 24 h with either DMSO or epigenetic compounds at indicated doses. Data are associated with Fig. 3 and 4.

**Supplementary Dataset 7.** Baseline and treatment-induced changes in population-averaged protein measurements of melanoma differentiation state markers (NGFR, AXL, MITF, SOX10), MAPK signaling protein modifications (p-$ERK^{T202/Y204}$, p-$S6^{S235/S236}$), a proliferation marker (Ki-67), and a DNA damage response marker (p-$H2A.X^{S139}$), across eight melanoma cell lines following treatment with epigenetic compounds, either individually or in combination with vemurafenib, or vemurafenib plus trametinib. Cells were pre-treated with the indicated epigenetic compounds for 24 h and then treated for a period of 3-5 days with either DMSO, vemurafenib (at 100 nM) or the combination of vemurafenib (at 100 nM) and trametinib (at 10 nM). Data are associated with Fig. 5a.

**Supplementary Dataset 8.** Baseline and treatment-induced changes in single-cell protein measurements of NGFR, AXL and MITF across eight melanoma cell lines following treatment with epigenetic compounds, either individually or in combination with vemurafenib, or vemurafenib plus trametinib. Cells were pre-treated with the indicated epigenetic compounds for 24 h and then treated for a period of 3-5 days with either DMSO, vemurafenib (at 100 nM) or the combination of vemurafenib (at 100 nM) and trametinib (at 10 nM). Single-cell data for a maximum of 1000 randomly selected cells per condition are shown. Data are associated with Fig. 5a.

**Supplementary Dataset 9.** Baseline and treatment-induced changes in single-cell protein measurements of p-ERK$^{T202/Y204}$, p-S6$^{S235/S236}$ and Ki-67 across eight melanoma cell lines following treatment with epigenetic compounds, either individually or in combination with vemurafenib, or vemurafenib plus trametinib. Cells were pre-treated with the indicated epigenetic compounds for 24 h and then treated for a period of 3-5 days with either DMSO, vemurafenib (at 100 nM) or the combination of vemurafenib (at 100 nM) and trametinib (at 10 nM). Single-cell data for a maximum of 1000 randomly selected cells per condition are shown. Data are associated with Fig. 5a.

**Supplementary Dataset 10.** Baseline and treatment-induced changes in single-cell protein measurements of SOX10 and p-H2A.X$^{S139}$ across eight melanoma cell lines following treatment with epigenetic compounds, either individually or in combination with vemurafenib, or vemurafenib plus trametinib. Cells were pre-treated with the indicated epigenetic compounds for 24 h and then treated for a period of 3-5 days with either DMSO, vemurafenib (at 100 nM) or the combination of vemurafenib (at 100 nM) and trametinib (at 10 nM). Single-cell data for a maximum of 1000 randomly selected cells per condition are shown. Data are associated with Fig. 5a.

**Supplementary Dataset 11.** Measurements of live cell count following exposure of the additional 5 melanoma cell lines (A101D, HS695T, IGR39, RPMI7951, SKMEL5) to either DMSO, vemurafenib (at 100 nM), or vemurafenib (at 100 nM) plus trametinib (at 10 nM) for a period of 72 or 120 h following pretreatment for 24 h with either DMSO, JIB-04 or SP2509 at indicated doses. Data are associated with Fig. 6d.

**Supplementary Dataset 12.** Baseline multiplexed single-cell measurements of NGFR and AXL proteins across 21 melanoma cell lines. Data are associated with Fig. 6d.

**Supplementary Dataset 13.** Baseline single-cell measurements of KDM4B protein across 21 melanoma cell lines. Data are associated with Fig. 6d.

**Supplementary Dataset 14.** Baseline single-cell measurements of ZNF217 protein across 21 melanoma cell lines. Data are associated with Fig. 6d.
